# Vocal coordination and conflict avoidance shape fission-fusion dynamics in white-nosed coatis

**DOI:** 10.64898/2026.05.26.727854

**Authors:** Emily M. Grout, Odd T. Jacobson, Jack C. Winans, Gabriella E. C. Gall, Josué Ortega, Julian C. Schaefer-Zimmermann, Margaret C. Crofoot, Ben T. Hirsch, Ariana Strandburg-Peshkin

## Abstract

Many social animals exhibit fission-fusion dynamics, where group members split into subgroups and come back together. Studying these dynamics can reveal how individuals weigh the costs and benefits of sociality. Most studies infer the drivers of fission-fusion dynamics by examining subgroup association patterns, yet different underlying processes can produce similar patterns. Here we take a fundamentally different approach by focusing on subgrouping events themselves, where decision-making plays out in real time. Using multi-sensor tracking collars, we collected simultaneous movement and vocalisation data from all group members within two wild white-nosed coati groups. We developed analytical tools to extract and characterise fission and fusion events, thus establishing a framework for quantifying the spatiotemporal dynamics of these events and the use of vocalisations before, during, and after them. We found that fissions typically occurred when groups were initially stationary, indicating that splitting does not result from loss of coordination while moving.Subgrouping was associated with reduced aggression, yet aggressive vocalisations did not precede splits, supporting the hypothesis that subgrouping is a pre-emptive mechanism to manage within-group conflict. Contact calls in moving subgroups increased before fissions and fusions, indicating that vocal communication is key to the coordination of these collective movement decisions.

## Introduction

Fission-fusion dynamics describe socio-spatial patterns in which the size and composition of animal groups change over time as individuals split into smaller subgroups and/or merge together (Aureli et al. 2008). These subgrouping patterns allow individuals to dynamically balance the costs (e.g. foraging competition) and benefits (e.g. reduced predation risk) of group living. Previous studies investigating fission-fusion dynamics have examined the temporal and spatial patterns of subgroup formations (Archie et al. 2006; Smith et al. 2008; Carter et al. 2013; Ramos-Fernández and Morales 2014; Aguilar-Melo et al. 2018; Della Libera et al. 2023), as well as their social and ecological drivers (Chapman et al. 1995; Smith et al. 2008; Ramos-Fernández and Morales 2014; Silk et al. 2014; Bond et al. 2019; Aguilar-Melo et al. 2020; Matthews et al. 2021; Hartwell et al. 2021). The collective decision-making processes underlying fissions and fusions themselves are still not well understood, largely because of the difficulties in gathering data from all group members across these events.Advances in tracking technology can provide a “birds-eye view” of group movement patterns, offering new perspectives that direct observation cannot achieve. By collecting data on the locations and behaviours of all group members simultaneously across fissions and fusions, we can explore the mechanisms by which these events occur, gaining insights into the proximate drivers of these dynamics.

White-nosed coatis, *Nasua narica*, form stable social groups in which membership remains largely constant, but these groups sometimes break up into smaller foraging parties and come back together (Kaufmann 1962; Gompper 1997; McColgin et al. 2018; Grout et al. 2024). Groups are typically composed of philopatric adult females and their offspring, while adult males are generally solitary, only joining groups during the mating season (Kaufmann 1962; Gompper and Krinsley 1992). Coatis produce a variety of vocalisations which provide information on their behavioural state and motivation, and which are believed to mediate group cohesion and social relationships (Kaufmann 1962; Gilbert 1973; Maurello et al. 2000; Compton et al. 2001; Hass 2021). Previous research found that when splitting into subgroups, coatis associate with close relatives rather than preferentially associating with others based on sex or age (Grout et al. 2024). Yet, the behavioural mechanisms driving fission and fusion events remain unknown. Using simultaneous movement and vocalisation data from all members of wild white-nosed coati groups, we investigate how and why fissions and fusions occur. To do so, we develop a generalisable method to identify and characterise the spatiotemporal properties of fission and fusion events, then use this method to test competing hypotheses regarding the drivers of fission and fusion dynamics in white-nosed coatis.

There are several hypothesised phenomena which could cause groups to split (Table S1; Figure 1F):1) Differences in activity budgets among group members, potentially driven by variation in body size or locomotor capacity, could lead to varying travel speed preferences, causing subgroups with different preferred speeds to separate from one another due to difficulties in maintaining cohesion (Ruckstuhl 1998; Conradt and Roper 2000; Ruckstuhl and Neuhaus 2002; Pontzer and Wrangham 2006; Sankey et al. 2019; Harel et al. 2021). If this is the case, we would expect splits to occur more often while a group is travelling with no change in individuals’ calling behaviour throughout these events. 2) Different foraging preferences could lead to conflicts of interest within a group over when and where to travel. If a conflict over where to travel arises, we would expect the group to split either when stationary or when moving, with the resultant subgroups heading in different directions (King et al. 2008; Sueur et al. 2011). In contrast, a conflict over when to travel would be expected to occur when a group is stationary and one subgroup decides to leave, or when a group is moving and one subgroup decides to stop (Papageorgiou and Farine 2020; Davis et al. 2022). As we would expect these movement decisions to be coordinated via vocal communication, we would expect that the group members that change their movement behaviour should produce contact calls more frequently throughout these events. 3) Foraging competition could drive groups to split to reduce the risk of receiving within-group aggression (Asensio et al. 2008). If so, we would expect splits to occur primarily when a group is stationary with one subgroup departing to forage elsewhere, or when a group is moving and one subgroup diverges to a different area. We would also expect higher rates of aggression when the group is together compared to when in subgroups, with aggression escalating in the period immediately preceding a fission. For hypotheses 2 and 3, we would expect group members to emit contact calls more frequently when they change their movement behaviour, as we hypothesise that they should use vocal communication to coordinate with one another throughout these events.These hypotheses are not mutually exclusive; fissions could be driven by a combination of these factors, or different events could have different underlying causes.

**Figure 1.**
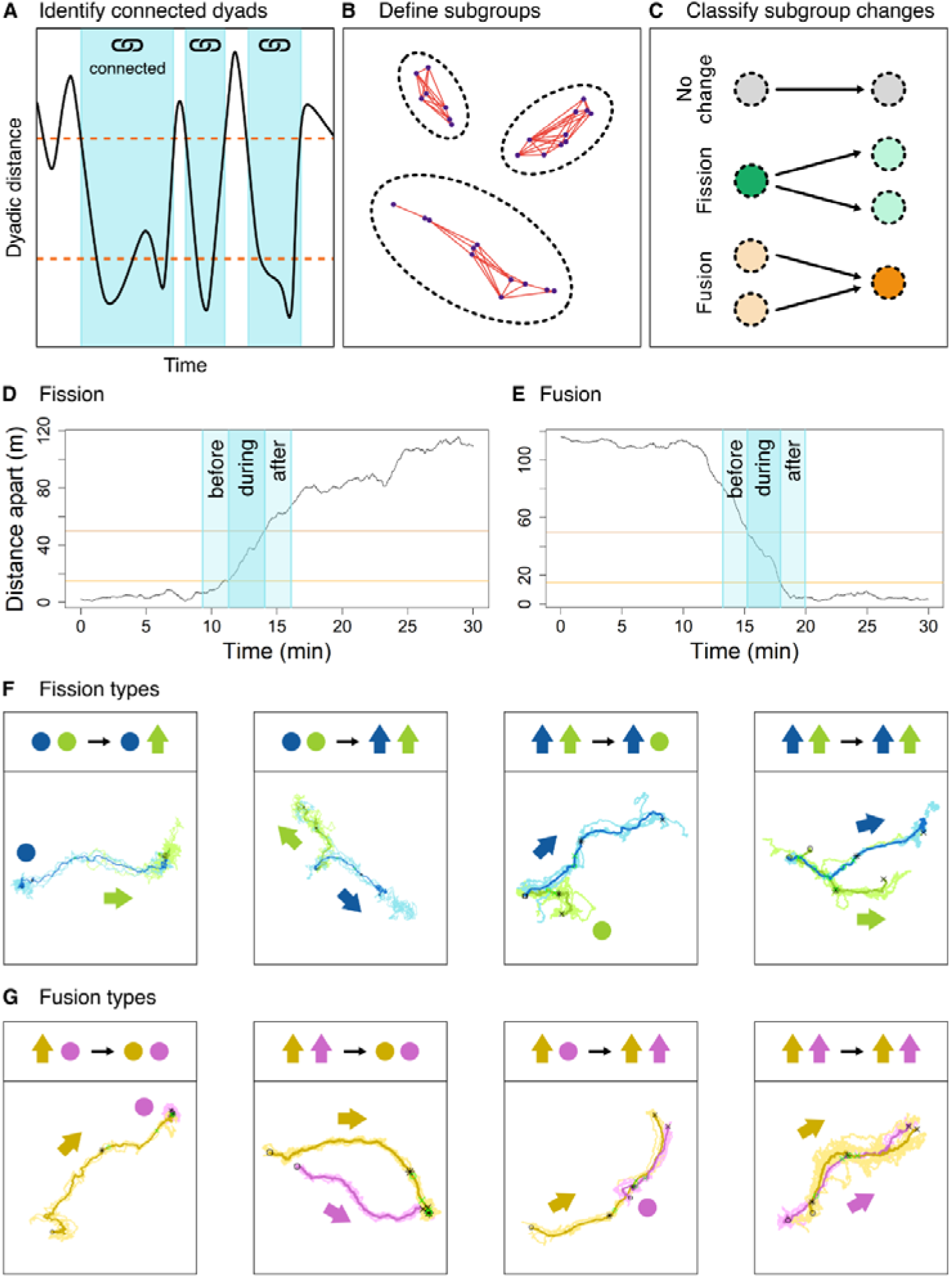
Illustration of how fission-fusion events were identified (A-C) and characterized (D-G) from tracking data. (A) Illustration for how connected periods between pairs of individuals (shaded in blue) are identified using a double threshold method based on dyadic distances between the individuals. (B) Subgroup membership is defined from periods of connectedness. Points represent the location of group members, red edges represent the nodes that are connected to one another based on the double threshold method, and dotted blue circles highlight subgroups that share members using the chain rule. (C) Bipartite graph showing the four possible options for changes in group membership between consecutive time points. Points represent groups identified in each time step and arrows link groups that share members. (D) Distance between the centroids of subgroups which fissioned. (E) Distance between the centroids of subgroups which fusioned. Vertical blue lines in (D) and (E) indicate the before time, start time, end time, and after time of these events, and horizontal orange lines indicate the 15 m and 50 m thresholds used for extracting these time points of interest. (F, G) Four types of fissions (F) and fusions (G), defined based on whether subgroups were stationary or moving *before* and *during* the event.

Group fusions can be shaped by several different factors (Table S1): 1) Subgroups may fuse as a result of attraction to the same resource (Cortés-Avizanda et al. 2011; Bennitt et al. 2018). If so, we expect that different parties will move towards the same location, or one subgroup will move towards another subgroup that is already at a desired destination. After a fusion, we expect groups to be stationary; and aggression call rates may be higher due to increased foraging competition (Aureli and Schaffner 2007). If calls are used to coordinate travel during fusions, we would expect a subgroup that moves to join a stationary subgroup to have an elevated contact call rate, whereas the stationary group being joined should show no change in contact call rate throughout the fusion event. 2) Fusions could occur because group members are attracted to one another, as observed in African elephants, *Loxodonta africana* (Archie et al. 2006), and orangutans, *Pongo pygmaeus* (Van Schaik 1999). If this is the case, we would either expect to observe the same movement patterns predicted above, or that subgroups travel together after merging. We would also expect both subgroups to have a higher contact calling rate throughout a fusion event to assist with locating one another (Spehar and Di Fiore 2013). Once the subgroups have merged, we would expect aggression levels to increase due to higher competition. As with fissions, these hypotheses regarding the drivers of group fusions are not mutually exclusive, and for many species, fusions are likely driven by both factors (Smith et al. 2008; Bond et al. 2019; Aguilar-Melo et al. 2020; Matthews et al. 2021) (Figure 1G).

To disentangle the different drivers of fissions and fusions in white-nosed coatis, we use multi-sensor tracking collars to simultaneously record the movements and vocalisations of all members of two wild social groups with different demographic structures, allowing us to assess the generality of the behavioural patterns observed. Using both movement and vocal data, we examine the behavioural mechanisms underlying fission-fusion dynamics, providing insight into how these group-living mammals manage the costs and benefits of sociality.

## Methods

To investigate the processes underlying fission and fusion events, we employed a multi-step methodological approach. First, we identified the occurrence of these events. We then characterised their distinct phases, which were used to extract movement-related properties. Finally, we integrated vocal behavioural data across these phases to assess the role of signalling in relation to these events. Each component of the methodology is described in detail below. Figures were prepared using RStudio (R Core Team 2017) and refined in Inkscape (Version 1.3.2).

### Study site and data collection

We conducted the study in the lowland tropical forests of Soberania National Park (9° 12’ N, -79° 70’ W) and on Barro Colorado Island (9° 16’ N, -79° 83’ W), Panama. All research activities were approved by MiAmbiente (No.SE/a-38-2020) and received animal ethics clearance from the STRI IACUC (2017-0815-2020). We equipped two groups of wild white-nosed coatis with custom built collars which housed a GPS sensor (e-Obs Digital Telemetry, Gruenwald, Germany) and two small microphones per collar (Soroka 18E, TS-Market) to record the vocalisations and positions of group members simultaneously. We also collared a third group, but, as they did not exhibit subgrouping behaviours, we did not include them in further analyses (see Grout *et al*., 2024 for further details). We caught coatis using Tomahawk traps and chemically immobilized them using Telazol (50 mg/ml tiletamine and 50 mg/ml zolazepam; 5.4 ± 0.5 mg/kg). Capture and collaring methods followed the ethical guidelines set by the American Society of Mammologists (see Grout *et al*., 2024 for further details on capture and collar deployment methods).

We recorded tracking data from the first group (Galaxy) for 17 days and the second group (Presidente) for 15 days. The groups differed in their demographic and relatedness structures. The Galaxy group consisted of 11 individuals (seven adult females, three subadults, and one juvenile), whereas the Presidente group consisted of 16 individuals (three adult females, seven subadults, and six juveniles). Each day from 0600 to 0900 EST, the GPS unit recorded one fix per second. The mean GPS fix success was > 96.1% for all collars, and the relative GPS error between two collars at known distances apart was 3.86 ± 1.06 m (mean ± standard deviation).

We recorded audio data from 0600 to 0900 to correspond with the 1Hz GPS period. Audio was recorded at 24000 Hz with 0 dB gain and 16-bit resolution. Due to the battery consumption of the audio recorders, each collar housed two devices to maximise the duration of audio recording, with the second recorder programmed to start 7 to 8 days after the first recorder (each recorder lasted 8.12 ± 0.7 days).

### GPS data processing

To examine the fine-scale movement patterns across fission and fusion events, we used GPS data from the 1Hz periods, which we pre-processed to remove fixes in unrealistic locations and to exclude periods of abnormal or missing data. Specifically, we excluded a time period when one individual (Venus) from the Galaxy group did not display normal behaviours, as this coati remained stationary in a tree for three days after being collared. Collars worn by two individuals (Moscoso and Peron) in the Presidente group fell off prematurely, and these individuals were subsequently recaptured and their collars were replaced. We removed the data collected in the periods when the collars were not on these animals, resulting in a gap of 5 hours and 101 hours for Moscoso and Peron, respectively.Moscoso’s collar fell off outside the 1 Hz sampling period, and therefore did not affect the high-frequency sampling data used in this study. We used RStudio (R Core Team 2017) for all pre-processing and quantitative analyses.

### Audio data processing

The audio recorders experienced an internal clock drift of approximately 8 seconds per day. To ensure synchronisation with the GPS data, we emitted synchronisation sounds during the recording period, which were picked up on the audio devices. The real times of these synchronisation sounds were used to accurately align the audio time with the GPS time in post-processing. Coati calls were defined using previous descriptions of the white-nosed coati vocal repertoire. Calls used in affiliative contexts include the *chirp grunt, chirp click, click grunt, click*, and *chirp* (Kaufmann 1962; Gilbert 1973; Smith 1977; Maurello et al. 2000; Compton et al. 2001; Hass 2021). Calls used in aggressive contexts include the *chitter* and *squeal* (Kaufmann 1962; Gilbert 1973; Smith 1977; Compton et al. 2001; Hass 2021). We manually labelled 62346 calls from a subset of audio data from all individuals in the Galaxy group using Adobe Audition (Adobe Inc. 2024). For subsequent automated call classification, we adapted the recently developed animal2vec model (Schäfer-Zimmermann et al. 2025), a large-scale transformer model (Vaswani et al. 2017) tailored for sparse multi-label bioacoustics data. Its training scheme consists of a self-supervised pre-training phase (Tarvainen and Valpola 2017; Baevski et al. 2022) on unlabelled data, followed by supervised fine-tuning on the labelled data. After finetuning, we performed inference on all available audio files from the Galaxy and Presidente group. Precision and Recall for the *squeal* class were too low for us to include them in our analysis; we therefore discarded them. Only *chitter* calls were considered as aggression calls. As *squeals* are often emitted after *chitter* calls during aggressive interactions, removing these calls is unlikely to affect our results, since we measured the presence or absence of aggression calls rather than call rates (see the Call Activity Analysis). Based on an initial training/validation split of the labelled data set and subsequent manual verification of detected calls, we estimated that contact calls were detected with a precision of 0.81 and a recall of 0.81, leading to 102373 and 137358 contact calls for the Galaxy and Presidente groups, respectively. Aggression calls were detected with a precision of 0.76 and a recall of 0.91, leading to 11426 and 10249 aggression calls for the Galaxy and Presidente groups, respectively (for further details, see ‘Audio Data Processing’ in Supplementary Methods; Figure S1).

### Detecting fissions and fusions

We detected fission and fusion events from simultaneous tracking data using a two-step procedure (top row in Figure 1) in which we first determined which individuals were in which subgroup at every moment in time and then compared subgroup membership over consecutive time steps to identify moments when subgroups split (fissions) or merged (fusions).

To determine subgroup membership every second, we used a modified version of the density-based spatial clustering algorithm (DBSCAN) (Ester et al. 1996), known as “sticky DBSCAN” (Della Libera et al. 2023). This algorithm used two dyadic distance thresholds (an inner and outer radius). The inner radius was used to detect instances in which two individuals were considered to be together (connected), whereas the outer radius was used to detect the beginning and end points of time periods of being connected. This double threshold increased the stability of periods by preventing rapid switching between connected and disconnected states that would occur if a single distance threshold were used (Della Libera et al. 2023). Missing data were handled by ending a period of connectedness (if present) whenever an individual stopped being tracked - in other words individuals were not considered to be connected with any other individuals when they had missing data. We identified subgroups by building up groups of individuals that were connected to at least one other subgroup member, similar to the “chain rule” often employed in observational field studies (Ramos-Fernández 2005). In other words, if individual i was connected to individual j and individual j was connected to individual k, then all three individuals (i, j, and k) would be considered to be in the same subgroup,even if i and k were not connected. Note that if an individual was not connected to any others, it was considered as a subgroup of size one.

Once we had identified subgroups at each moment in time, we examined changes in subgroup membership across time to identify fissions and fusions. To do so, we first created a bipartite graph where the first set of nodes (set A) represented the subgroups present at the current time (t) and the second set of nodes (set B) represented the subgroups present at the next time point (t + 1). A node from set A was then considered linked to a node from set B if the groups contained any of the same members. There were four possible patterns arising from the bipartite graphs: (1) a single node from set A linked to a single node from set B (no change in subgroup membership), (2) a single node from set A linked to multiple nodes from set B (a *fission*), (3) multiple nodes from set A linked to a single node from set B (a *fusion*), and (4) multiple nodes from set A linked to multiple nodes from set B (a *shuffle*). Using this approach, we extracted fissions and fusions, as well as identified how many subgroups were involved and which individuals were in each subgroup. In the current dataset, no shuffles and no fissions or fusions involving more than 2 subgroups were observed. Because we were examining the social and communicative aspects of splitting and merging events, fissions and fusions in which a single individual joined or left a group were excluded from subsequent analyses (Figure S2; Galaxy: 25 fissions, 33 fusions; Presidente: 9 fissions, 8 fusions).

### Characterising event phases (before, during, and after)

Although the method described above identified a single time point at which subgroup composition changed, in reality, fissions and fusions occurred over an extended window of time as the two (or more) subgroups moved apart or came together. To characterise fissions and fusions, we first identified a *start time* and an *end time* for each event, with the time window between these two points considered as the period over which the event took place (row 2 in Figure 1). Our method was designed to handle events (fissions and fusions) involving only two subgroups, as we did not observe any events involving three or more subgroups in our data.

To identify start and end times, we first computed the centroids (mean x-y position) of the two subgroups associated with the event at every moment in time around the initially-detected event time (*t_event*) within a time window of *t_event –* 700 s to *t_event* + 700 s. For a fission, these were the subgroups that resulted from the split, whereas for a fusion, these were the two subgroups that came together during the merge. We then defined the start and end of events using two thresholds, *D*_*lower*_ and *D*_*upper*_. For fissions, we identified contiguous sequences where the distance between the two centroids (*D*_*centr*_) went from below the lower threshold (*D*_*centr*_ *< D*_*lower*_) to between the two thresholds (*D*_*lower*_ *< D*_*centr*_ *< D*_*upper*_) to above the upper threshold (*D*_*centr*_ *> D*_*upper*_) over a period of time. The *start time* of the event was identified as the time when the distance between the two centroids crossed the lower threshold (*D*_*lower*_); this represented the time at which the two subgroups were close together. The *end time* of the event was defined as the time when the distance between the two centroids rose above the upper threshold (*D*_*upper*_); this represented the time at which the two subgroups had separated substantially from one another. If multiple sequences were identified during the time window around an event, the times closest to the initially identified event time were used. For fusions, the same method was used but in reverse; the *start time* was identified as when the distance between centroids was above *D*_*upper*_ and the *end time* was identified as when the distance between centroids dropped below *D*_*lower*_. We set *D*_*lower*_ to 15 m and *D*_*upper*_ to 50 m, with these thresholds chosen based on visual inspection of the data. Due to the multi-scale nature of these events, and because we were approximating subgroup locations with the centroids, the dyadic distance sometimes did not reach below the lower threshold or above the upper threshold. In these cases, we modified the upper or lower threshold to the maximum or minimum dyadic distance before and after the initially identified event time within the 700 s time window.

In addition to the start and end time of each event, we also identified a *before time* and *after time* to represent the behaviour of the group immediately preceding or following the event. Here, the *before time* was defined as the time point 120 s before the *start time* and the *after time* was defined as the time point 120 s after the *end time*. If the *before* or *after* time of an event would run into a preceding or subsequent event, the *before* or *after* time was defined as the time point immediately preceding the start or end of the adjacent event. Using this method, we identified three phases for each event; *before* the event (before time to start time), *during* the event (start time to end time), and *after* the event (end time to after time). Methods for identifying and characterizing fissions and fusions are illustrated in Figure 1.

### Characterising event properties

To determine how fissions and fusions occurred, we characterised these events based on whether each of the two subgroups were *moving* or *stationary* during different phases of the event, allowing us to categorise each event into an event *type* (see Figure 1 F,G). Based on this categorisation scheme, a fission could occur in four distinct ways (Figure 1F): (i) when a single subgroup moved off from an initially stationary subgroup, (ii) when both subgroups moved off in different directions from an initially stationary subgroup, (iii) in an initially moving group, when one subgroup stopped moving and the other continued, or (iv) in an initially moving group, where both subgroups continued moving but in different directions. Fusions events could also be similarly categorised (Figure 1G). To categorise whether subgroups were moving or stationary during each phase, we used their spatial displacement during each phase. For fissions, the displacements were computed using the centroid of all members in the group *before* the fission (from before time to start time), and the centroid of each subgroup *during* the event (from start time to end time). For fusions, the displacements were computed using the centroid of each subgroup *during* the event (from start time to end time), and the centroid of all members in the group *after* the fusion (from end time to after time). The displacement was defined as the distance between the centroid location at the beginning and end of the specified period. We then classified whether a subgroup was moving or stationary *during* fissions and fusions using a threshold. Based on field observations when groups were stationary, and by visual inspection of the distribution of distance travelled *during* fission and fusion events, we chose a 10m displacement of the subgroup centroid as the threshold for classifying whether a subgroup was moving or stationary *during* the event (see Figure S3 for distributions of distance travelled).

### Call activity analysis

We investigated the role vocal communication played with respect to fission and fusion events in coatis using calls classified by the machine learning model. For each individual, we extracted contact call and aggression call activity for the entire study duration as well as during fission and fusion events. For fission and fusion events, we quantified call activity within the *before, during*, and *after* periods, as defined above. The duration of the ‘*during*’ period varied across events as it was determined by the time taken to meet the spatial thresholds defining each event (Presidente: 147 ± 73 s; Galaxy: 191 ± 106 s; Figure S4). The *before* and *after* periods each spanned two minutes immediately preceding or following the event. Call rates around fission and fusion events were compared to baseline periods (when groups were together or in subgroups but not in the vicinity of a fission or fusion event), quantified over two-minute intervals. For contact calls, we used the total count of calls per individual within each period. For aggression calls, because they are rare but tend to occur in rapid bursts when they do occur, raw counts would overrepresent their prevalence. We therefore used a binary indicator of the presence or absence of aggression calls per individual within each period.

### Travel speed

We calculated travel speed by down-sampling the GPS data to 30-second intervals to minimise error from GPS noise. We then extracted the mean speed for each group member in two-minute bins for periods when the group was together. We excluded data from when fewer than 9/11 or 14/16 individuals were tracked in the Galaxy group and the Presidente group respectively.

### Statistical Analyses

Our statistical analysis aimed to understand how vocal communication related to fission-fusion dynamics. Because our predictions concerned different aspects of these dynamics (subgroup size, the temporal phases of fission-fusion events, and the movement roles of subgroups) we used separate generalised linear mixed-effects models (GLMMs) to address each prediction, all fitted using the brms package (Bürkner 2021). In all models, the unit of analysis was the individual: for contact calls, the response variable was the number of calls produced by an individual within a given time period, modelled with a zero-inflated negative binomial distribution; for aggression calls, the response was the binary presence or absence of aggression calls per individual (see Call Activity Analysis for details), modelled with a Bernoulli distribution. All models included varying intercepts per individual to account for repeated observations.

To test the prediction that call activity differs between subgroup and full-group contexts, we included subgroup size as a categorical predictor indicating whether the individual was in a subgroup (4–6 and 3–13 individuals for Galaxy and Presidente, respectively) or a full group (>8 and >13 individuals for Galaxy and Presidente, respectively). We used a categorical rather than continuous predictor because the distribution of group sizes was bimodal, with observations clustered at typical subgroup and full-group sizes and few intermediate values (Figure S5), a continuous predictor was poorly suited to capture this relationship. Because aggression calls are typically louder than contact calls, we included median dyadic distance as an additional covariate in the aggression call model to control for sound “leakage” between devices (i.e., calls from one individual being recorded on nearby individuals’ devices). This metric was calculated for each focal individual, in each 2-minute bin, as the median of its distances to all subgroup members.

To evaluate how call activity varies as a function of travel speed independent of fission and fusion events, we fit models with travel speed (standardised across both groups), group identity, and their interaction as fixed effects, restricting the data to bouts in which the full group was together (as defined above). The random-effects and distributional structures were as described above, with varying intercepts per individual. As a sensitivity check, we refit each model with varying intercepts and slopes for speed per individual; population-level inferences were qualitatively unchanged, and the per-individual fits are shown in the supplementary material (Figure S6).

To test the prediction that call activity changes across the temporal phases of fission and fusion events, we modelled fissions and fusions separately, with the fission-fusion period (*together, before, during, after*) as a categorical predictor. The ‘*together*’ category served as a baseline (“control”) representing periods when the full group was together and not near a fission or fusion event. These models included an exposure variable in the linear predictor to account for variation in the duration of event periods. To test the prediction that moving and stationary subgroups differ in their call activity *during* events, we fitted additional models with the subgroup’s movement role as a predictor. Fissions and fusions were modelled separately: in fission models, subgroup roles were categorised as leavers (departing) or remainers (stationary); in fusion models, subgroup roles were categorised as joiners (approaching) or stationed (stationary). This model focused on the most common event type, in which one subgroup was stationary and one was moving (Figure 2A), and included an interaction between the fission-fusion period and subgroup role. Baseline comparisons were excluded from this model.

**Figure 2.**
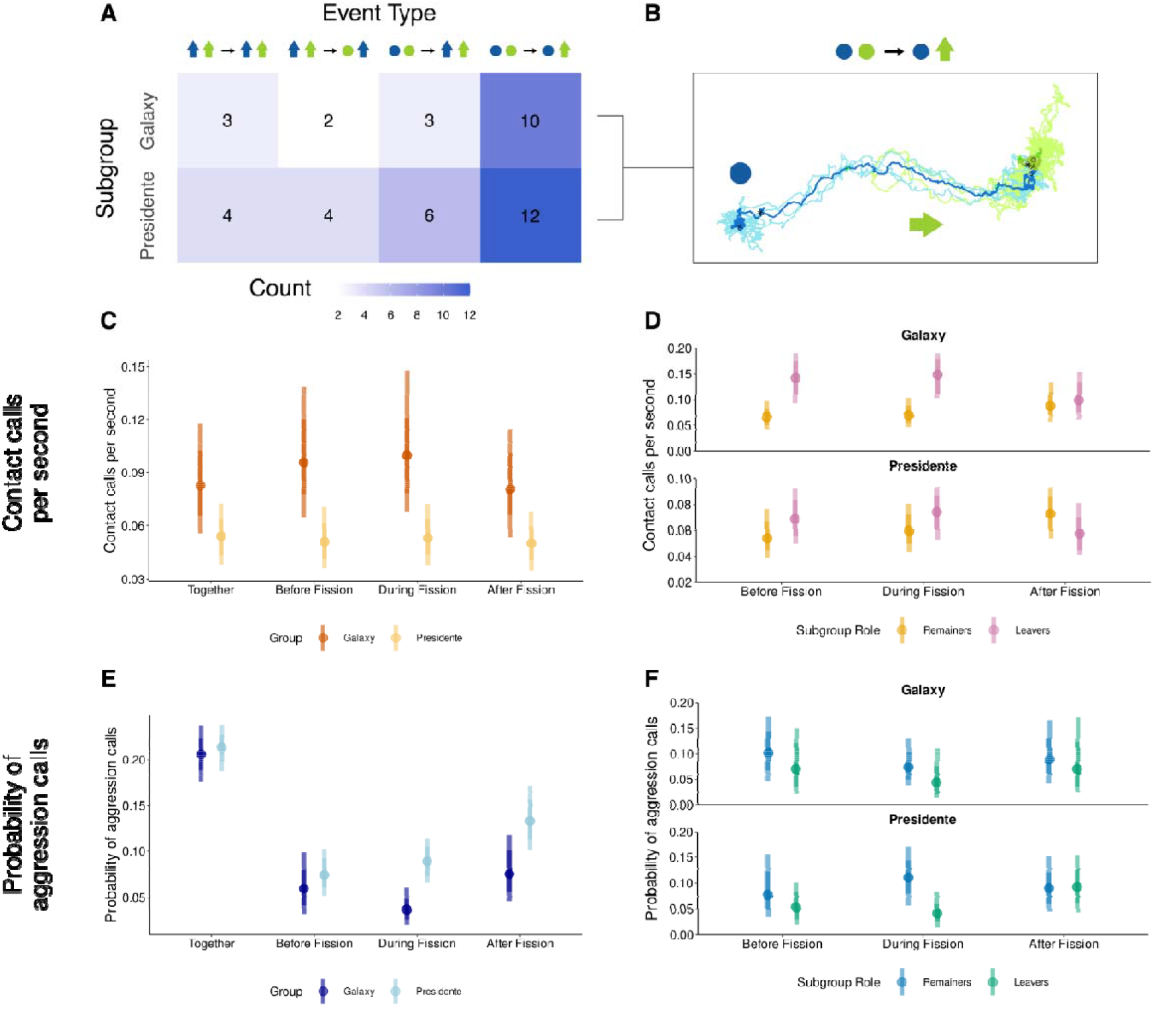
Spatial dynamics and calling activity associated with group fissions. (A) Count of each fission event type observed in the Galaxy group (left) and the Presidente group (right). (B) Trajectory example of the most frequent fission event type, where a group is initially stationary and one subgroup splits off while the other remains stationary. (C) Posterior point intervals showing the model-estimated median along with 66% and 89% credible intervals for contact call rate across different fission periods, including baseline comparisons (when the group was together) for the Galaxy group and the Presidente group. (D) Model predictions for contact call rate based on the interaction between fission period and subgroup action (leaving vs remaining) for the Galaxy group and the Presidente group. Orange represents the subgroup that remained stationary and did not initiate the fission event (remainers) and pink represents the subgroup that moved away *during* the fission event (leavers). (E) Posterior point intervals showing the model-estimated median along with 66% and 89% credible intervals for aggression call probability across different fission periods, including baseline comparisons (when the group was together) for the Galaxy group and the Presidente group. (F) Model predictions for aggression call probability based on the interaction between fission period and subgroup action (leaving vs remaining) for the Galaxy group and the Presidente group. Blue represents the subgroup that remained stationary and did not initiate the fission event (remainers) and green represents the subgroup that moved away *during* the fission event (leavers). Note that the axes differ between the two plots in D and F.

To assess whether non-independence between individuals within the same event affected our results, we conducted a sensitivity analysis in which we refitted the fission-fusion period models with an additional varying intercept per event. Because baseline observations are not associated with any event and therefore cannot accommodate this random effect, we compared models with and without the event-level intercept using only the event periods (before, during, after). The inclusion of the event-level random intercept did not substantively change the magnitude or direction of the estimated effects (Figure S7), indicating that our primary results are robust to this source of non-independence.

We assessed model fit using posterior predictive checks, comparing observed data with simulated outcomes from each model’s posterior predictive distribution. Continuous predictors were z-score standardised to facilitate comparison and improve model fit. The effects of categorical predictors were evaluated by computing posterior differences between levels. We report the proportion of the posterior greater than zero (PP > 0) and 89% Highest Posterior Density Intervals (HPDI) to characterise the credibility and uncertainty of parameter estimates.

## Results

### Fission events are driven by subgroup departure and coordinated via vocal communication

Most fissions occurred while the group was stationary; one subgroup departed while the other remained in place (Figure 2), rather than fissioning during group travel (as would have been expected under the activity budget hypothesis, Hypothesis 1; see Table S1 for details). When groups were together, faster travel was associated with higher contact call rates (Figure S8; Galaxy: β = 0.259 [0.219, 0.303]; Proportion of the posterior greater than 0 (PP > 0): 1; Presidente: β = 0.089 [0.061, 0.121]; PP > 0: 1), suggesting that contact calls play a role in coordinating group movement.Consistent with this, individuals in leaving subgroups had higher contact call rates before and during fission events than individuals in remaining subgroups, with no difference after the split (Figure 2D; Galaxy: β(during) _leavers − remainers_ = 0.831 [0.438, 1.21]; PP > 0: 1; Presidente: β(during) _leavers − remainers_ = 0.242 [-0.002, 0.489]; PP > 0: 0.939). Together, these results suggest that departing individuals use contact calls to coordinate their movement away from the group during fission events.

### Aggression is higher in full groups but does not trigger fission events

The probability of aggression calls per individual was higher when in full groups compared to subgroups (Figure 3B, c; Galaxy: β = 1.082 [0.863, 1.314]; PP > 0: 1; Presidente: β = 0.439 [0.169, 0.684]; PP > 0: 0.995), and did not vary with travel speed when groups were together (Figure S8; Galaxy: β = -0.037 [-0.194, 0.137]; PP > 0: 0.35; Presidente: β = -0.003 [-0.104, 0.106]; PP > 0:0.475). The probability of aggression calls was also lower during all fission phases (before, during, after) compared to other times when full groups were together (Figure 8b; Galaxy: β_during fission - together_ = -1.894 [-2.405, -1.35]; PP > 0: 0; Presidente: β_during fission - together_ = -1.021 [-1.332, -0.726]; PP > 0: 0), indicating that heightened aggression does not immediately precede splits.

**Figure 3.**
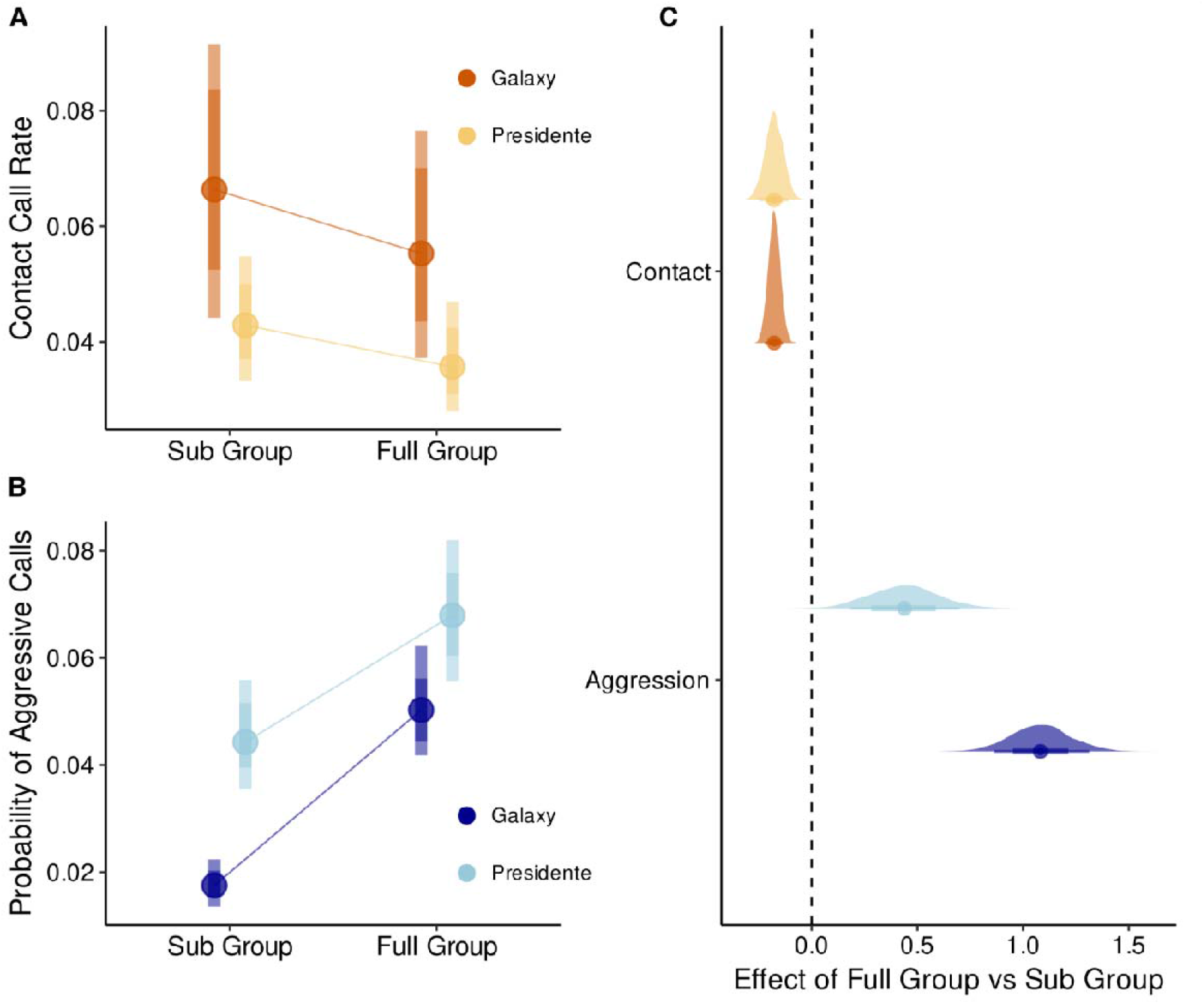
Call rates vary as a function of group context. Model predictions for the effect of sub vs. full group indicator on contact call rates (orange) and probability of aggression calls (blue) for the Galaxy group (dark shade) and the Presidente group (light shade). (A, B) Posterior point intervals showing the median and 66% and 95% credible intervals. (C) Posterior densities and 50% and 95% credible intervals for the effect size coefficients of the sub/full group indicator. Note that a positive effect size indicates a stronger effect of full groups compared to subgroups and vice versa.

### Vocal coordination within joining subgroups facilitates fusion events

Most fusions involved one subgroup (joiners) moving towards a stationary subgroup (Figure 4), consistent with both the resource attraction and social attraction hypotheses. For these events, contact call rates were higher for joiners than for the stationary subgroup before and during fusion events (Figure 4D; Galaxy: β_(before) joiners – stationed_ = 1.011 [0.697, 1.322]; PP > 0: 1; Presidente: β_(before) joiners – stationed_ = 0.791 [0.552, 1.003]; PP > 0: 1; Galaxy: β_(during) joiners – stationed_ = 0.377 [0.083, 0.674]; PP > 0:0.980; Presidente: β_(during) joiners – stationed_ = 0.498 [0.274, 0.732]; PP > 0: 1). Contact call rates for stationary subgroups increased across fusion events (Galaxy: β_(stationed)after – before_ = 0.743 [0.485, 1.036]; PP > 0: 1; Presidente: β_(stationed)after – before_ = 0.147 [-0.043, 0.321]; PP > 0: 0.898), indicating that the stationary subgroup increased calling as the joiners approached.

**Figure 4.**
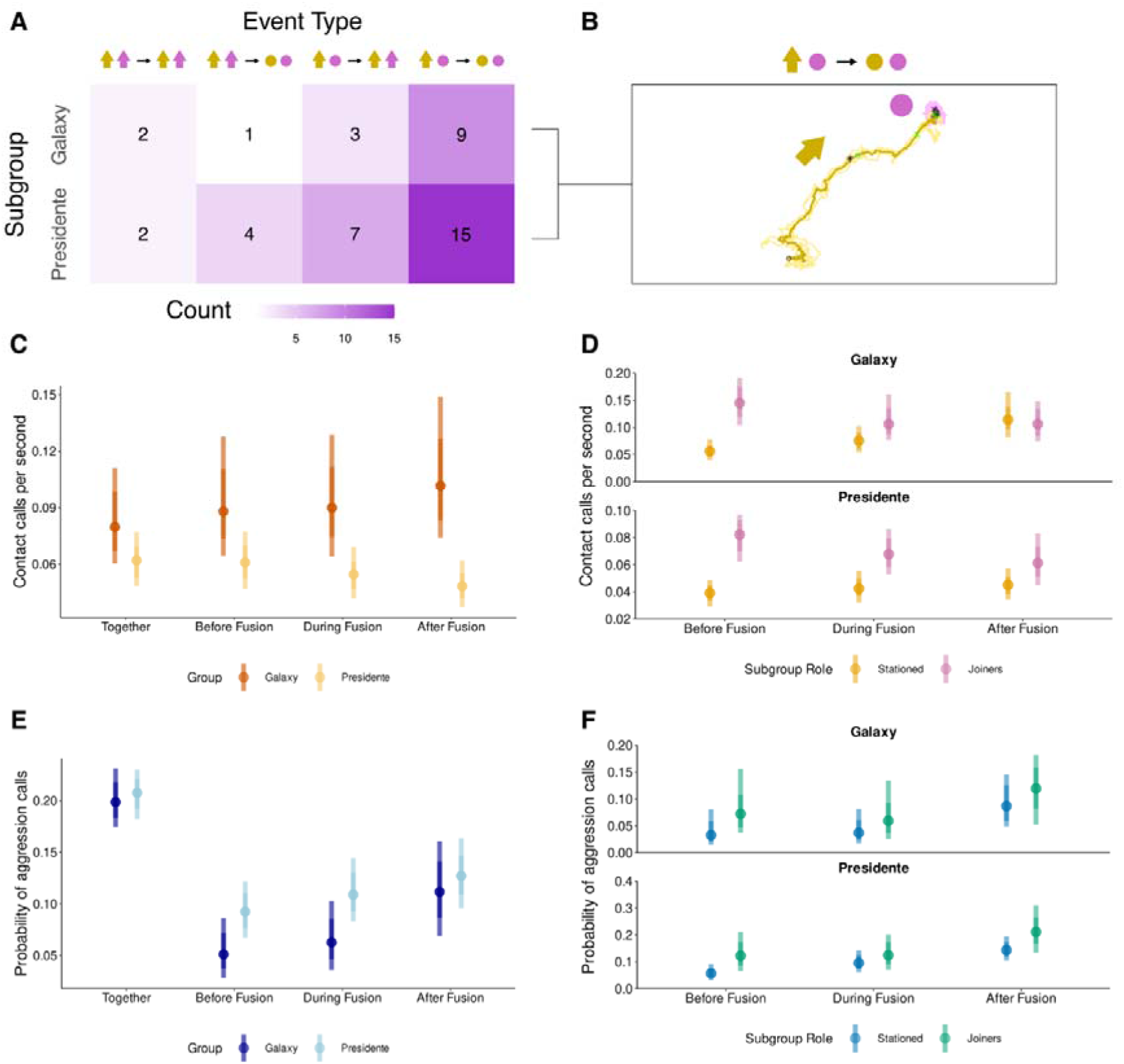
Spatial dynamics and calling activity associated with group fusions. (A) Count of each fusion event type observed in the Galaxy group (left) and the Presidente group (right). (B) Trajectory example of the most frequent fusion event type, where a moving subgroup joins an initially stationary subgroup. (C) Posterior point intervals showing the model-estimated median along with 66% and 89% credible intervals for contact call rate across different fusion periods, including baseline comparisons. (D) Model predictions for contact call rate based on the interaction between fusion period and subgroup action (joining vs stationed). Orange represents subgroups that remained stationary and did not initiate the fusion event (stationed). Pink represents the subgroup that moved toward the stationary subgroup *during* the fusion event (joiners). (E) Posterior point intervals showing the model-estimated median along with 66% and 89% credible intervals for aggression call probability across different fusion periods, including baseline comparisons. (F) Model predictions for aggression call probability based on the interaction between fusion period and subgroup action (joining vs stationed). Blue represents subgroups that remained stationary and did not initiate the fusion event (stationed) and green represents the subgroup that moved toward the stationary subgroup *during* the fusion event (joiners). Note that the axes differ between the two plots in D and F.

### Aggression decreases around fusion events

The probability of aggression calls was lower before, during, and after fusion events compared to the baseline in both groups (Figure 4E; Galaxy: β_(during fusion − together)_ = -1.331 [-1.809, -0.777]; PP > 0: 0; Presidente: β_(during fusion − together)_ = -0.746 [-1.023, -0.431]; PP > 0: 0). There was a slight increase in the probability of aggression calls from before to after fusion events (Galaxy: β_(after - before)_ = 0.826 [0.113, 1.568]; PP > 0: 0.967; Presidente: β_(after - before)_ = 0.356 [-0.039, 0.783]; PP > 0: 0.910). Though there was an overall reduction in aggression around fusion events, the increase in the probability of aggression after a fusion is consistent with the resource attraction hypothesis, which predicts increased foraging competition upon merging.

## Discussion

Our results suggest that fission events are driven by conflicts over the timing of departure decisions and may be a strategy to avoid aggression, while fusions are driven by attraction to group members and/or shared resources. Vocal communication plays a central role in coordinating both processes, with contact calls facilitating subgroup departure during fissions and enabling approaching subgroups to coordinate their movements during fusions. Crucially, aggression does not trigger splits, but is higher when groups are together, suggesting that subgrouping may serve as a pre-emptive strategy to manage within-group conflict.

By simultaneously recording the movements and vocalisations of all group members, we reveal the behavioural mechanisms underlying these collective decisions, a perspective unattainable through direct observation alone. This study demonstrates that fission-fusion dynamics in white-nosed coatis are driven by conflicts of interest within groups over the timing of behavioural state-changes, rather than differences in preferred travel speeds. By focusing on behavioural processes, rather than their emergent patterns, we show that most fissions occurred while groups were stationary, with one subgroup departing, suggesting that splits are driven by conflicting preferences on *when* to travel, and do not arrive as a consequence of loss of coordination during travel itself. Two plausible explanations could account for this pattern: (a) groups may have been resting prior to a split, with a fission occurring due to conflicts of interest over when to resume foraging; or (b) groups may have been foraging at a clumped resource such as a fruit tree or a dense invertebrate patch (Kaufmann 1962; Gompper 1995; Valenzuela 1998; Hirsch 2009; Hirsch et al. 2024), with one subgroup choosing to leave the patch earlier than the other. Future efforts to disentangle these possibilities would require behavioural observations of group members’ activity states before fission events, and it is also possible that both types of events occur.

Although coatis are known to engage in aggressive competition at clumped resources (Kaufmann 1962; Gompper 1996; Gompper et al. 1997; Hirsch 2007, 2011; De La O et al. 2019), we found no evidence that it served as a proximate cause of group splits in our data. In fact, we found a lower probability of aggression calls before splits compared to baseline periods when the full group was together, contrasting with other fission-fusion species such as spotted hyenas where aggression has been observed to directly precede fission events (Smith et al. 2008). Rather than aggressive interactions leading to group splits, our results point to the *avoidance* of conflict as an underlying motivation for splitting. Aggression was higher overall when groups were together than when in subgroups. This is consistent with previous findings that coati groups split along relatedness lines (Grout et al. 2024), likely as a mechanism to avoid conflict during direct foraging competition, given that unrelated individuals receive a disproportionate amount of aggression (Gompper et al. 1997).Together, these results suggest that group members split pre-emptively to reduce the risk of aggression from particular individuals, rather than in direct response to aggressive encounters.

While both attraction to shared resources and attraction to the other subgroup may explain fusions, our results provide stronger support for the latter. In both study groups, the majority of fusions involved a moving subgroup joining a stationary subgroup, after which they remained stationary together. The probability of aggression increased across fusion events and was slightly higher in the joining subgroup compared to the stationary subgroup, suggesting that changes in the social environment increased tension within the group. Individuals in the joining subgroup emitted contact calls at a higher rate before and during these events, consistent with using vocal communication to coordinate their approach. Calling rates also increased in the stationary subgroup, suggesting potential vocal signalling between subgroups during the merge, consistent with the hypothesis that fusions are driven by attraction to the other subgroup. How the moving subgroup locates the stationary subgroup remains unclear, particularly given that they inhabit dense forest where subgroups may travel more than 100 m apart. Possible mechanisms include eavesdropping on vocalisations produced by the stationary subgroup, using olfactory cues, or navigating toward known sites (e.g. high-quality foraging or resting areas) rather than toward the other subgroup itself (Strauss et al. 2024). As coatis are a highly olfactory species (Chapman 1938; Compton 1973; Gompper 1996; Hirsch 2010), with recently released captured individuals observed to follow scent trails of their group members (personal observation), it is likely olfaction plays a large role in locating the other subgroup. Because aggression calls are considerably louder than contact calls, they may be detectable at greater distances, raising the possibility that moving subgroups could eavesdrop on these calls to locate stationary subgroups. To disentangle the sensory processes governing fusions, future studies should compare the movement paths of individuals searching for group mates with the paths previously travelled by those group mates, as well as determining the range at which coatis can detect conspecific vocalisations, in order to assess the relative importance of following scent trails versus relying on auditory cues.

Overall, our results suggest that a combination of social and ecological factors shape fission-fusion dynamics in white-nosed coatis. Although aggression probabilities are higher when groups are together, coatis gain several benefits from being in a larger group, including reduced predation risk and resource defence against adult males (Gompper 1996; Hass and Valenzuela 2002; Hirsch and Gompper 2018). The fact that subgroups consistently coalesce back into larger groups suggests that, despite increased aggression in the full group, the benefits of group living ultimately outweigh the costs, with fission-fusion dynamics providing a flexible strategy for managing this trade-off.Moreover, some fusions involved a moving subgroup joining a stationary subgroup after which they continued travelling together, suggesting that a shared resource was not the sole reason for coming back together. Determining what behaviours group members are engaging in before, during, and after fissions and fusions would give further insight into the drivers of these events.

The finding that group splits occur primarily when groups are stationary suggests that these events reflect differences in patch departure times, which has important implications for social foraging and consensus decision-making. A substantial body of research has investigated ecological drivers of patch departure decisions such as predation risk and resource availability (Brown 1988, 1999; Bowers et al. 1993; Druce et al. 2006; Rieucau et al. 2009; Arehart et al. 2023). For group-living animals, these decisions are also socially contingent, as maintaining cohesion often requires individuals to reach consensus on when to leave a foraging patch (Giraldeau and Caraco 2000; Davis et al. 2022).When optimal departure timing varies substantially among group members, individuals may either compromise with the full group or split into subgroups. Fission-fusion dynamics may therefore allow animals such as coatis to relax the constraint of group-wide consensus, minimising consensus costs while maximising individual foraging success (Kerth 2010; Sueur et al. 2011). Our finding that contact calls increase specifically in the departing subgroup suggests that vocal communication provides the mechanism through which such partial consensus is reached. Because our movement data were collected during the first three hours of daily activity, when coatis are motivated to forage after the overnight fast (Kaufmann 1962), stationary periods before fissions most likely represent foraging bouts rather than rest periods, strengthening the interpretation that splits are linked to patch departure decisions.

While splitting into subgroups offers a potential mechanism to reduce consensus costs, these decisions are further complicated by social preferences. Coatis typically split based on relatedness, with subgroups comprising individuals of different age and sex classes (Grout et al. 2024), meaning that optimal departure times likely still vary among individuals within each subgroup. Decision-making conflicts therefore likely persist even after subgroup formation, suggesting that coatis navigate a complex landscape in which they must balance maximising individual foraging success, maintaining cohesion with preferred social partners, and avoiding individuals from whom they are more likely to receive aggression (Gompper et al. 1997). Despite these residual conflicts, splitting reduces the number of competitors at clumped resources, enhancing foraging success in the longer term (Gompper 1996; Hirsch 2009). Examining intake rates of group members before fission events would help clarify how individuals balance their own foraging needs with maintaining cohesion with preferred social partners (Gompper 1996). Because coatis are far from unique in displaying both fission-fusion dynamics and consistent social preferences, these insights are likely to have broad implications for understanding the dynamics of collective decision-making in group-living species.

Our study provides a generalisable framework to characterise different fission and fusion types and to quantify the role of vocalisations during these processes, providing a new perspective on subgrouping dynamics. Recent advances in tracking technology have enabled detailed inference of the drivers of collective behaviour in cohesive social groups (Biro et al. 2006; Nagy et al. 2010; King et al. 2012; Strandburg-Peshkin et al. 2015; Farine et al. 2017), and our framework extends this approach to fission-fusion dynamics, where the mechanisms underlying collective decisions remain comparatively understudied. To build on these findings, future studies should investigate how the resource landscape, predation risk, and variation in social organisation shape the spatial and vocal dynamics of fission-fusion events across species and ecological contexts. Integrating tracking data with behavioural observations during fission and fusion events would further resolve how social and ecological pressures interact to drive collective behaviour.

## Author Contributions

EMG: Conceptualisation, Data curation, Formal analysis, Investigation, Methodology, Project administration, Visualisation, Software, Writing – original draft; OTJ: Formal analysis, Investigation, Methodology, Visualisation, Writing – review & editing; JW: Code Review, Methodology, Writing – review & editing; GECG: Investigation, Methodology, Writing – review & editing; JO: Data curation, Project administration; JSZ: Data curation, Methodology, Software, Writing – review & editing; MCC: Conceptualization, Funding acquisition, Investigation, Project administration, Supervision, Writing – review & editing; BTH: Conceptualization, Data curation, Funding acquisition, Investigation, Methodology, Project administration, Supervision, Writing – review & editing; ASP: Conceptualization, Data curation, Funding acquisition, Investigation, Methodology, Project administration, Software, Supervision, Writing – review & editing

## Acknowledgements

We thank the Smithsonian Tropical Research Institute, MiAmbiente, and the Republic of Panama for permission to conduct this research. We are grateful to Rachel Page, Melissa Cano, and Lil Camacho at the Smithsonian Tropical Research Institute for their support during fieldwork. We thank Patrick Paetzold for designing and customising the collar casing which held audio recorders. Thank you to Carolina Mitre Ramos, Brandol Ortega, and Lucía Torrez for fieldwork assistance, as well as Julia Plecher, Leonie Treyer, Caroline Preuß, and Roshni Keshwani, and Zeynep Durmus for their assistance in labelling and validating the audio data. We thank Christine Hass for sharing her knowledge on coati vocalisations. We are grateful to all members of the Ecology of Animal Societies Department, Communication and Collective Movement Research Group, and Communication and Coordination Across Scales Project for discussion and feedback. We especially thank Imran Razik for assistance with figure visualisations and Alison Ashbury for writing support. Funding supporting this research was provided by a Human Frontier Science Program Research Grant (RGP0051/2019) to A.S.P. and B.T.H., the Max Planck Society, the Gips-Schüle Stiftung, and the Centre for the Advanced Study of Collective Behaviour (EXC 2117–422037984).

