## Supplementary materials for "Vocal coordination and conflict avoidance shape fission-fusion dynamics in white-nosed coatis"

Table S1. Hypotheses and predictions for fission and fusion events, with a qualitative assessment of whether each prediction is supported by the results (full, partial, or not supported).

| Type | Hypothesis | Predicted movement patterns | Predicted calling patterns | Support |
| --- | --- | --- | --- | --- |
| Fission | Different activity budgets | The group is travelling | No change in calling behaviour | None |
| Fission | Different preferences over *where* to go | The group is stationary and resultant subgroups travel to different locations.  or  The group is moving and resultant subgroups travel in different directions. | Group members that change their movement behaviour emit contact calls more frequently. | Partial |
|  | Different preferences over *when* to go | The group is stationary and one subgroup leaves.  or  The group is moving and one subgroup stops. | Group members that change their movement behaviour emit contact calls more frequently. | Full |
| Fission | Foraging competition | The group is stationary and one subgroup leaves.  or  The group is moving and subgroups go to different areas. | Greater aggression when the group is together compared to when in subgroups.  Probability of aggression is greater before a fission.  Group members that change their movement behaviour emit contact calls more frequently. | Partial |
| Fusion | Attraction to the same resource | Subgroups move towards the same location  or  One subgroup moves towards another subgroup that is already at the destination.  After a fusion, group will be stationary. | Greater aggression when the group is together compared to when in subgroups.  Group members that change their movement behaviour emit contact calls more frequently.  Stationary group members will not change contact call rate. | Partial |
| Fusion | Attraction to one another | Subgroups move towards the same location  or  One subgroup moves towards another subgroup that is already at the destination.  After a fusion, group will be stationary.  or  Group travel together after merging. | Group members that change their movement behaviour emit contact calls more frequently.  Stationary group members will emit contact calls more frequently (to facilitate reunion).  Greater aggression when the group is together compared to when in subgroups. | Full |

*Audio Data Processing*

The audio-processing procedure used in this study was also applied in (Winans et al. 2026), and is therefore also described in that study’s supplementary materials.

The audio recorders generated a large data set of raw audio files (> 1,800 h). To identify and classify coati vocalizations from this data set, we trained a self-supervised transformer-based neural network model (animal2vec, (Schäfer-Zimmermann et al. 2025)) on 115760 manually-labelled vocalizations that were annotated by trained analysts from a subset of audio files in Adobe Audition (Adobe Inc. 2024). Label files consisted of contiguous sequences of audio for which the onsets and offsets of all calls within that segment were labelled. Each audio label encoded two classifications: (i) call type and (ii) focal status. The focal status of each call could either be classified as ‘focal’ (meaning the call was thought to be given by the individual wearing the collar) or ‘nonfocal’ (meaning the call was thought to be given by another individual within detection range of the recorder mounted on the focal individual’s collar). Human labellers primarily distinguished between ‘focal’ and ‘nonfocal’ calls based on call amplitude and additional contextual information where available, but we note that these classifications are likely prone to greater error than classifications for call type. animal2vec was trained and tested using all 10-sec audio clips for which at least one acoustic event was present, following Schäfer-Zimmermann et al. 2025. A subset of the manually-labelled calls (n = 28970 calls) were excluded from the training set to serve as a validation set. With this validation set, we achieved an average precision of 0.847 micro averaged across call types (Zhu 2004).

After training animal2vec to detect coati calls and classify them by call type and focal status, we used the trained model to generate call detections (onset time, offset time, call type, and focal status) across all audio files in our data set. The animal2vec model typically produces predictions on a fine-grained, frame-by-frame basis; here, we adopt the post-processing method described by the developers (Schäfer-Zimmermann et al. 2025). Event (i.e. call) boundaries are determined by applying a sliding average-pooling window to the likelihood output, which is subsequently binarized through a fixed threshold. The likelihood score for the segment is then the mean of the frame-wise likelihoods within the predicted sequence. This workflow effectively converts frame-level data into instance-level metrics, resulting in a single prediction for each discrete event. The likelihood per event is bounded between 0 and 1, which indicated the model’s confidence that the call belonged to the assigned class. For each likelihood score, we assessed the recall (fraction of real calls of a given class that were correctly detected as that class) and precision (fraction of detected calls of a given class that actually belonged to that class). To assess the recall, we used the subset of data held out from training. However, because this hold-out set only included 10-second clips in which at least one acoustic event was detected, computing the precision based on this set would likely result in an underestimate, because a large proportion of the raw audio files consists of audio for which no calls are present but the machine learning model might still produce false detections. Therefore, to get an accurate assessment of precision, we generated predictions from animal2vec across 25 complete (3-hour) audio files that were previously not used in training, and that represented all individuals in the dataset (one file per individual), then manually verified the call type and focal status of the resulting 45,954 machine learning-generated detections. Using these verified files, we then computed the precision associated with each likelihood score based on how many detections were correct and how many were false positives. Finally, we combined these values with the associated recall score at each likelihood value to generate the precision-recall curves shown in Figure S1. For the purposes of this study, we were specifically interested in call types hypothesised to serve as ‘contact calls’: *chirp*, *chirp click*, *chirp grunt*, *click*, and *click grunt*. Therefore, we collapsed these acoustically distinguishable calls into one hypothesised functional category hereafter referred to as ‘contact calls’, and calculated the precision-recall curve for this “contact call superclass”. Here, for each likelihood score, we defined the associated recall as the fraction of all (ground truth labelled) focal calls of any contact call type that were detected by animal2vec as any contact call type (using the hold-out set). We defined the precision associated with a given likelihood score as the fraction of detections of any focal contact call type that truly belonged to any focal contact call category (using the verified files).

The precision-recall curves and likelihood scores for the contact call superclass and the *chitter* class are shown in Figure S1. Based on these results, we chose a likelihood threshold of 0.4 for contact calls and a likelihood threshold of 0.1 for *chitters*. Using these thresholds, we estimate that contact calls were detected with a precision of 0.81 and a recall of 0.81 and *chitters* were detected with a precision 0.76 and a recall of 0.91 (see Figure S1).

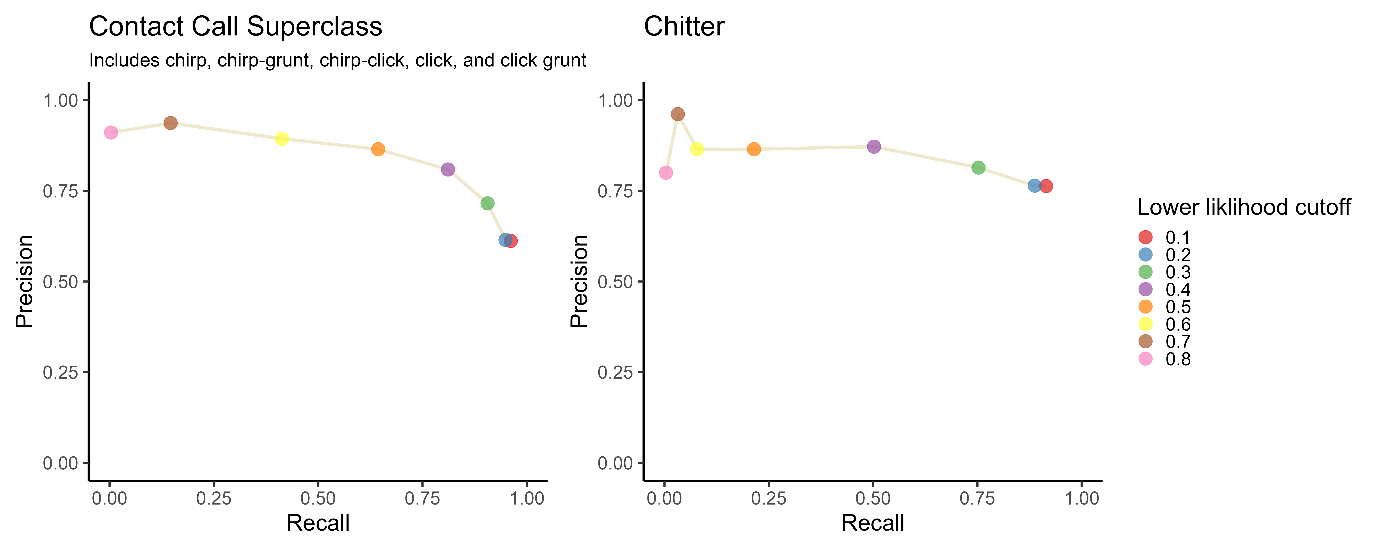

Figure S1. Estimated precision and recall of detected contact calls (left) and chitters (right) as a function of likelihood score. Likelihood scores were calculated for each machine learning-detected coati call, and each point represents the estimated precision and recall of call detections assigned a likelihood score greater than a given cut-off. Based on these estimates, we only analysed detected contact calls that were assigned a likelihood score of > 0.40 (purple point in left panel) and detected chitters that were assigned a likelihood score of > 0.10 (red point in right panel). These thresholds were chosen to optimally balance precision and recall. They led to a recall of 0.81 (81% of true calls detected) and a precision of 0.81 (81% of detected calls were correct) for contact calls, and a recall of 0.91 (91% of true calls detected) and a precision of 0.76 (76% of detected calls were correct) for chitters.

Table S2. List of response and predictor variables used in statistical models, as well as figures where results are presented.

| Response Variable |  | Predictors | Figure |
| --- | --- | --- | --- |
| Contact Call Rate | ~ | Mean Travel Speed | S8 |
| Aggression Call (0/1) | ~ | Mean Travel Speed | S8 |
| Contact Call Rate | ~ | Sub/Full Group Indicator | 3A |
| Aggression Call (0/1) | ~ | Sub/Full Group Indicator + Median Dyadic Distance | 3B |
| Contact Call Rate | ~ | Fission Period + Exposure (Bout Duration) | 2C |
| Aggression Call (0/1) | ~ | Fission Period + Exposure (Bout Duration) | 2E |
| Contact Call Rate | ~ | Fusion Period + Exposure (Bout Duration) | 4C |
| Aggression Call (0/1) | ~ | Fusion Period + Exposure (Bout Duration) | 4E |
| Contact Call Rate | ~ | Subgroup Action + Fission Period + Subgroup Action:Fission Period + Exposure (Bout Duration) | 2D |
| Aggression Call (0/1) | ~ | Subgroup Action + Fission Period + Subgroup Action:Fission Period + Exposure (Bout Duration) | 2F |
| Contact Call Rate | ~ | Subgroup Action + Fusion Period + Subgroup Action:Fusion Period + Exposure (Bout Duration) | 4D |
| Aggression Call (0/1) | ~ | Subgroup Action + Fusion Period + Subgroup Action:Fusion Period + Exposure (Bout Duration) | 4F |

Table S3. Posterior summaries for all model parameters, by group. Median = posterior median; P(>0) = proportion of posterior above zero; 89% HDI = 89% highest density interval. All estimates are on the link scale.

| Figure | Model | Group | Contrast | Median | P(>0) | 89% HDI |
| --- | --- | --- | --- | --- | --- | --- |
| S8 | Contact ~ Speed | Galaxy | Slope of speed (std) | 0.259 | 1.000 | [0.219, 0.303] |
|  |  | Presidente | Slope of speed (std) | 0.089 | 1.000 | [0.061, 0.121] |
| S8 | Aggression ~ Speed | Galaxy | Slope of speed (std) | -0.037 | 0.350 | [-0.194, 0.137] |
|  |  | Presidente | Slope of speed (std) | -0.003 | 0.475 | [-0.104, 0.106] |
| 3A | Contact ~ Subgroup Size | Galaxy | contrast full_group - sub_group | -0.177 | 0.000 | [-0.22, -0.132] |
|  |  | Presidente | contrast full_group - sub_group | -0.178 | 0.000 | [-0.244, -0.107] |
| 3B | Aggression ~ Subgroup Size | Both (Not Interacted) | Median dyadic distance (std) | -0.292 | 0.000 | [-0.385, -0.193] |
|  |  | Galaxy | contrast full_group - sub_group | 1.082 | 1.000 | [0.863, 1.314] |
|  |  | Presidente | contrast full_group - sub_group | 0.439 | 0.995 | [0.169, 0.684] |
| 2C | Contact ~ Fission Period (+ baseline) | Galaxy | contrast base - before | -0.157 | 0.016 | [-0.273, -0.022] |
|  |  | Galaxy | contrast base - during | -0.192 | 0.006 | [-0.309, -0.068] |
|  |  | Galaxy | contrast base - after | 0.014 | 0.565 | [-0.114, 0.139] |
|  |  | Galaxy | contrast before - during | -0.035 | 0.375 | [-0.196, 0.139] |
|  |  | Galaxy | contrast before - after | 0.171 | 0.935 | [0.001, 0.354] |
|  |  | Galaxy | contrast during - after | 0.205 | 0.973 | [0.04, 0.376] |
|  |  | Presidente | contrast base - before | 0.047 | 0.813 | [-0.034, 0.138] |
|  |  | Presidente | contrast base - during | 0.005 | 0.535 | [-0.075, 0.096] |
|  |  | Presidente | contrast base - after | 0.071 | 0.899 | [-0.018, 0.154] |
|  |  | Presidente | contrast before - during | -0.043 | 0.261 | [-0.153, 0.065] |
|  |  | Presidente | contrast before - after | 0.024 | 0.628 | [-0.08, 0.139] |
|  |  | Presidente | contrast during - after | 0.066 | 0.831 | [-0.043, 0.177] |
| 2D | Contact ~ Fission Period x Ind. Action | Galaxy | [Period \| remainers] contrast before - during | -0.074 | 0.349 | [-0.376, 0.251] |
|  |  | Galaxy | [Period \| remainers] contrast before - after | -0.330 | 0.044 | [-0.661, -0.035] |
|  |  | Galaxy | [Period \| remainers] contrast during - after | -0.260 | 0.090 | [-0.567, 0.049] |
|  |  | Galaxy | [Period \| leavers] contrast before - during | -0.031 | 0.453 | [-0.427, 0.367] |
|  |  | Galaxy | [Period \| leavers] contrast before - after | 0.444 | 0.955 | [0.031, 0.871] |
|  |  | Galaxy | [Period \| leavers] contrast during - after | 0.472 | 0.969 | [0.081, 0.886] |
|  |  | Galaxy | [Ind. Action \| before] contrast remainers - leavers | -0.879 | 0.000 | [-1.296, -0.494] |
|  |  | Galaxy | [Ind. Action \| during] contrast remainers - leavers | -0.831 | 0.000 | [-1.21, -0.438] |
|  |  | Galaxy | [Ind. Action \| after] contrast remainers - leavers | -0.098 | 0.351 | [-0.507, 0.32] |
|  |  | Presidente | [Period \| remainers] contrast before - during | -0.093 | 0.290 | [-0.364, 0.177] |
|  |  | Presidente | [Period \| remainers] contrast before - after | -0.330 | 0.026 | [-0.622, -0.072] |
|  |  | Presidente | [Period \| remainers] contrast during - after | -0.236 | 0.080 | [-0.49, 0.039] |
|  |  | Presidente | [Period \| leavers] contrast before - during | -0.088 | 0.261 | [-0.332, 0.124] |
|  |  | Presidente | [Period \| leavers] contrast before - after | 0.179 | 0.889 | [-0.054, 0.406] |
|  |  | Presidente | [Period \| leavers] contrast during - after | 0.266 | 0.972 | [0.051, 0.501] |
|  |  | Presidente | [Ind. Action \| before] contrast remainers - leavers | -0.246 | 0.066 | [-0.51, 0.007] |
|  |  | Presidente | [Ind. Action \| during] contrast remainers - leavers | -0.242 | 0.061 | [-0.489, 0.002] |
|  |  | Presidente | [Ind. Action \| after] contrast remainers - leavers | 0.262 | 0.950 | [0.008, 0.511] |
| 2E | Aggression ~ Fission Period (+ baseline) | Galaxy | contrast base - before | 1.401 | 1.000 | [0.887, 2.021] |
|  |  | Galaxy | contrast base - during | 1.894 | 1.000 | [1.35, 2.405] |
|  |  | Galaxy | contrast base - after | 1.128 | 1.000 | [0.616, 1.649] |
|  |  | Galaxy | contrast before - during | 0.500 | 0.859 | [-0.229, 1.224] |
|  |  | Galaxy | contrast before - after | -0.273 | 0.278 | [-1.014, 0.445] |
|  |  | Galaxy | contrast during - after | -0.768 | 0.045 | [-1.462, -0.039] |
|  |  | Presidente | contrast base - before | 1.217 | 1.000 | [0.826, 1.586] |
|  |  | Presidente | contrast base - during | 1.021 | 1.000 | [0.726, 1.332] |
|  |  | Presidente | contrast base - after | 0.566 | 0.999 | [0.283, 0.887] |
|  |  | Presidente | contrast before - during | -0.201 | 0.251 | [-0.667, 0.256] |
|  |  | Presidente | contrast before - after | -0.656 | 0.011 | [-1.128, -0.206] |
|  |  | Presidente | contrast during - after | -0.456 | 0.036 | [-0.873, -0.053] |
| 2F | Aggression ~ Fission Period x Ind. Action | Galaxy | [Period \| remainers] contrast before - during | 0.416 | 0.761 | [-0.497, 1.343] |
|  |  | Galaxy | [Period \| remainers] contrast before - after | 0.172 | 0.610 | [-0.806, 1.197] |
|  |  | Galaxy | [Period \| remainers] contrast during - after | -0.235 | 0.348 | [-1.175, 0.73] |
|  |  | Galaxy | [Period \| leavers] contrast before - during | 0.465 | 0.707 | [-0.945, 1.879] |
|  |  | Galaxy | [Period \| leavers] contrast before - after | -0.038 | 0.482 | [-1.524, 1.432] |
|  |  | Galaxy | [Period \| leavers] contrast during - after | -0.502 | 0.282 | [-1.892, 0.894] |
|  |  | Galaxy | [Ind. Action \| before] contrast remainers - leavers | 0.458 | 0.744 | [-0.645, 1.64] |
|  |  | Galaxy | [Ind. Action \| during] contrast remainers - leavers | 0.511 | 0.785 | [-0.538, 1.586] |
|  |  | Galaxy | [Ind. Action \| after] contrast remainers - leavers | 0.266 | 0.646 | [-0.847, 1.387] |
|  |  | Presidente | [Period \| remainers] contrast before - during | -0.412 | 0.248 | [-1.402, 0.52] |
|  |  | Presidente | [Period \| remainers] contrast before - after | -0.230 | 0.362 | [-1.303, 0.785] |
|  |  | Presidente | [Period \| remainers] contrast during - after | 0.183 | 0.628 | [-0.774, 1.071] |
|  |  | Presidente | [Period \| leavers] contrast before - during | 0.204 | 0.614 | [-0.926, 1.385] |
|  |  | Presidente | [Period \| leavers] contrast before - after | -0.684 | 0.152 | [-1.78, 0.385] |
|  |  | Presidente | [Period \| leavers] contrast during - after | -0.888 | 0.076 | [-1.939, 0.118] |
|  |  | Presidente | [Ind. Action \| before] contrast remainers - leavers | 0.511 | 0.775 | [-0.585, 1.612] |
|  |  | Presidente | [Ind. Action \| during] contrast remainers - leavers | 1.139 | 0.974 | [0.169, 2.115] |
|  |  | Presidente | [Ind. Action \| after] contrast remainers - leavers | 0.059 | 0.543 | [-0.882, 0.988] |
| 4C | Contact ~ Fusion Period (+ baseline) | Galaxy | contrast base - before | -0.107 | 0.104 | [-0.233, 0.029] |
|  |  | Galaxy | contrast base - during | -0.111 | 0.085 | [-0.244, 0.014] |
|  |  | Galaxy | contrast base - after | -0.243 | 0.002 | [-0.381, -0.12] |
|  |  | Galaxy | contrast before - during | -0.006 | 0.478 | [-0.182, 0.172] |
|  |  | Galaxy | contrast before - after | -0.139 | 0.118 | [-0.304, 0.05] |
|  |  | Galaxy | contrast during - after | -0.133 | 0.110 | [-0.317, 0.033] |
|  |  | Presidente | contrast base - before | 0.012 | 0.594 | [-0.069, 0.098] |
|  |  | Presidente | contrast base - during | 0.127 | 0.989 | [0.038, 0.214] |
|  |  | Presidente | contrast base - after | 0.243 | 1.000 | [0.16, 0.336] |
|  |  | Presidente | contrast before - during | 0.115 | 0.951 | [0.007, 0.227] |
|  |  | Presidente | contrast before - after | 0.231 | 1.000 | [0.117, 0.338] |
|  |  | Presidente | contrast during - after | 0.116 | 0.953 | [0.006, 0.232] |
| 4D | Contact ~ Fusion Period x Ind. Action | Galaxy | [Period \| stationed] contrast before - during | -0.292 | 0.040 | [-0.57, -0.026] |
|  |  | Galaxy | [Period \| stationed] contrast before - after | -0.743 | 0.000 | [-1.036, -0.485] |
|  |  | Galaxy | [Period \| stationed] contrast during - after | -0.451 | 0.004 | [-0.701, -0.181] |
|  |  | Galaxy | [Period \| joiners] contrast before - during | 0.338 | 0.949 | [-0.003, 0.66] |
|  |  | Galaxy | [Period \| joiners] contrast before - after | 0.375 | 0.960 | [0.035, 0.72] |
|  |  | Galaxy | [Period \| joiners] contrast during - after | 0.036 | 0.572 | [-0.312, 0.363] |
|  |  | Galaxy | [Ind. Action \| before] contrast stationed - joiners | -1.011 | 0.000 | [-1.322, -0.697] |
|  |  | Galaxy | [Ind. Action \| during] contrast stationed - joiners | -0.377 | 0.020 | [-0.674, -0.083] |
|  |  | Galaxy | [Ind. Action \| after] contrast stationed - joiners | 0.110 | 0.711 | [-0.214, 0.413] |
|  |  | Presidente | [Period \| stationed] contrast before - during | -0.088 | 0.222 | [-0.269, 0.098] |
|  |  | Presidente | [Period \| stationed] contrast before - after | -0.147 | 0.102 | [-0.321, 0.043] |
|  |  | Presidente | [Period \| stationed] contrast during - after | -0.060 | 0.300 | [-0.244, 0.121] |
|  |  | Presidente | [Period \| joiners] contrast before - during | 0.206 | 0.896 | [-0.053, 0.468] |
|  |  | Presidente | [Period \| joiners] contrast before - after | 0.339 | 0.981 | [0.071, 0.598] |
|  |  | Presidente | [Period \| joiners] contrast during - after | 0.133 | 0.795 | [-0.116, 0.405] |
|  |  | Presidente | [Ind. Action \| before] contrast stationed - joiners | -0.791 | 0.000 | [-1.003, -0.552] |
|  |  | Presidente | [Ind. Action \| during] contrast stationed - joiners | -0.498 | 0.000 | [-0.732, -0.274] |
|  |  | Presidente | [Ind. Action \| after] contrast stationed - joiners | -0.304 | 0.018 | [-0.544, -0.077] |
| 4E | Aggression ~ Fusion Period (+ baseline) | Galaxy | contrast base - before | 1.535 | 1.000 | [0.931, 2.147] |
|  |  | Galaxy | contrast base - during | 1.331 | 1.000 | [0.777, 1.809] |
|  |  | Galaxy | contrast base - after | 0.708 | 0.996 | [0.25, 1.173] |
|  |  | Galaxy | contrast before - during | -0.193 | 0.340 | [-1, 0.551] |
|  |  | Galaxy | contrast before - after | -0.826 | 0.033 | [-1.568, -0.113] |
|  |  | Galaxy | contrast during - after | -0.635 | 0.066 | [-1.304, 0.024] |
|  |  | Presidente | contrast base - before | 0.938 | 1.000 | [0.634, 1.279] |
|  |  | Presidente | contrast base - during | 0.746 | 1.000 | [0.431, 1.023] |
|  |  | Presidente | contrast base - after | 0.590 | 0.999 | [0.301, 0.888] |
|  |  | Presidente | contrast before - during | -0.188 | 0.228 | [-0.611, 0.218] |
|  |  | Presidente | contrast before - after | -0.356 | 0.090 | [-0.783, 0.039] |
|  |  | Presidente | contrast during - after | -0.159 | 0.254 | [-0.545, 0.236] |
| 4F | Aggression ~ Fusion Period x Ind. Action | Galaxy | [Period \| stationed] contrast before - during | -0.001 | 0.499 | [-1.058, 0.999] |
|  |  | Galaxy | [Period \| stationed] contrast before - after | -0.968 | 0.056 | [-1.907, 0.024] |
|  |  | Galaxy | [Period \| stationed] contrast during - after | -0.966 | 0.046 | [-1.904, -0.041] |
|  |  | Galaxy | [Period \| joiners] contrast before - during | 0.337 | 0.671 | [-0.851, 1.545] |
|  |  | Galaxy | [Period \| joiners] contrast before - after | -0.518 | 0.230 | [-1.608, 0.636] |
|  |  | Galaxy | [Period \| joiners] contrast during - after | -0.847 | 0.110 | [-1.955, 0.296] |
|  |  | Galaxy | [Ind. Action \| before] contrast stationed - joiners | -0.828 | 0.094 | [-1.881, 0.131] |
|  |  | Galaxy | [Ind. Action \| during] contrast stationed - joiners | -0.493 | 0.218 | [-1.474, 0.512] |
|  |  | Galaxy | [Ind. Action \| after] contrast stationed - joiners | -0.378 | 0.248 | [-1.228, 0.564] |
|  |  | Presidente | [Period \| stationed] contrast before - during | -0.564 | 0.106 | [-1.318, 0.141] |
|  |  | Presidente | [Period \| stationed] contrast before - after | -0.971 | 0.010 | [-1.686, -0.298] |
|  |  | Presidente | [Period \| stationed] contrast during - after | -0.406 | 0.141 | [-1.039, 0.192] |
|  |  | Presidente | [Period \| joiners] contrast before - during | 0.026 | 0.521 | [-0.841, 0.904] |
|  |  | Presidente | [Period \| joiners] contrast before - after | -0.558 | 0.137 | [-1.319, 0.285] |
|  |  | Presidente | [Period \| joiners] contrast during - after | -0.581 | 0.120 | [-1.46, 0.152] |
|  |  | Presidente | [Ind. Action \| before] contrast stationed - joiners | -0.849 | 0.051 | [-1.653, 0.013] |
|  |  | Presidente | [Ind. Action \| during] contrast stationed - joiners | -0.266 | 0.288 | [-1.034, 0.464] |
|  |  | Presidente | [Ind. Action \| after] contrast stationed - joiners | -0.434 | 0.145 | [-1.059, 0.254] |

All estimates are on the link scale (logit for Bernoulli/Beta Models, log for count/Gamma Models). Contrasts are computed via emmeans, conditioned on group using at = list(group = ...). For Models with additional covariates (e.g., median dyadic distance), emmeans holds these at their observed means. For Models with offset(log(duration)), offset is set to 0; since offsets cancel in pairwise contrasts on the link scale, contrast estimates are unaffected. P(>0) = proportion of the posterior distribution above zero. 89% HDI = 89% Highest Density Interval.

|  | Galaxy | Presidente |
| --- | --- | --- |
| Fission | 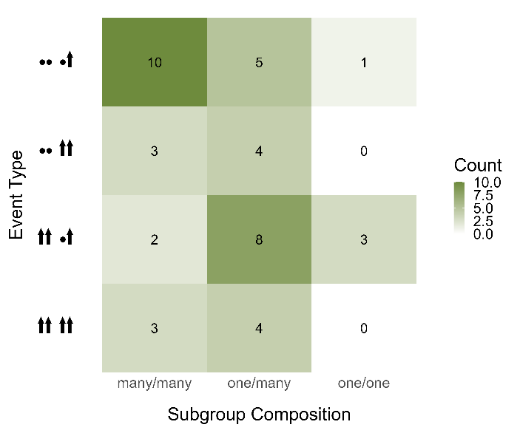 | 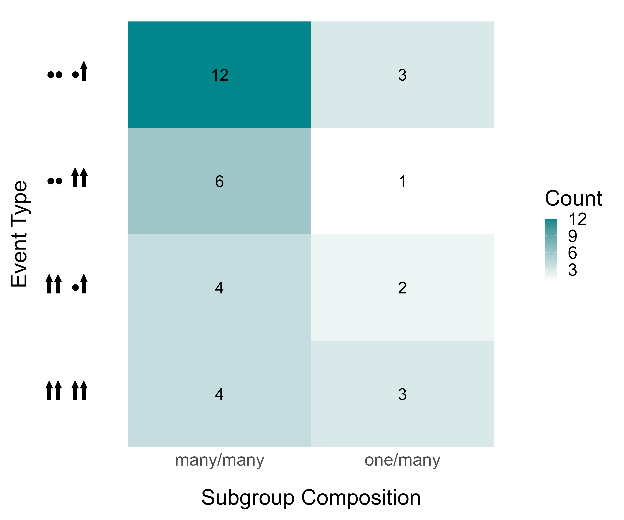 |
| Fusion | 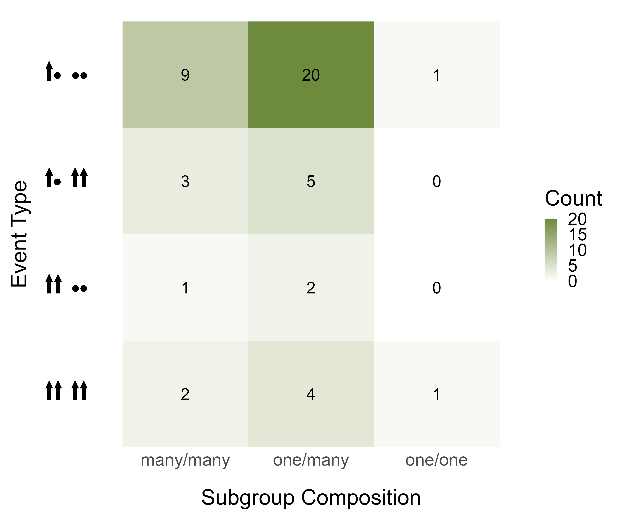 | 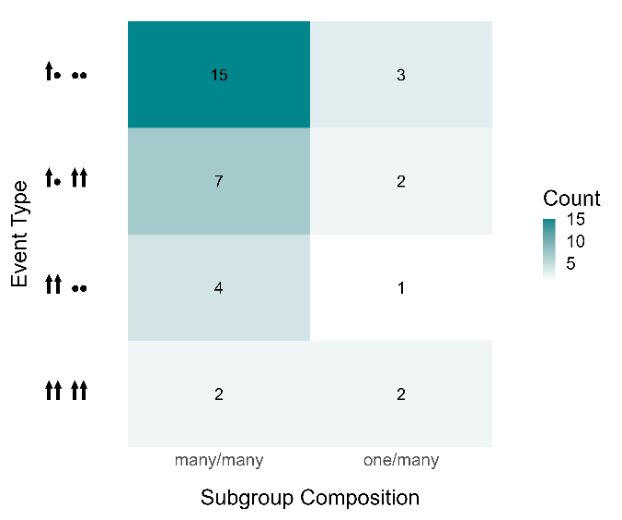 |

Figure S2. Counts of each fission (top row) and fusion (bottom row) type for different subgroup compositions in the Galaxy group (left) and the Presidente group (right), excluding events caused by males leaving the group. The compositions are categorised as follows: "many/many," where both subgroups have more than one individual; "one/many," where one subgroup has only one individual while the other has more than one; and "one/one," where each subgroup contains only one individual.

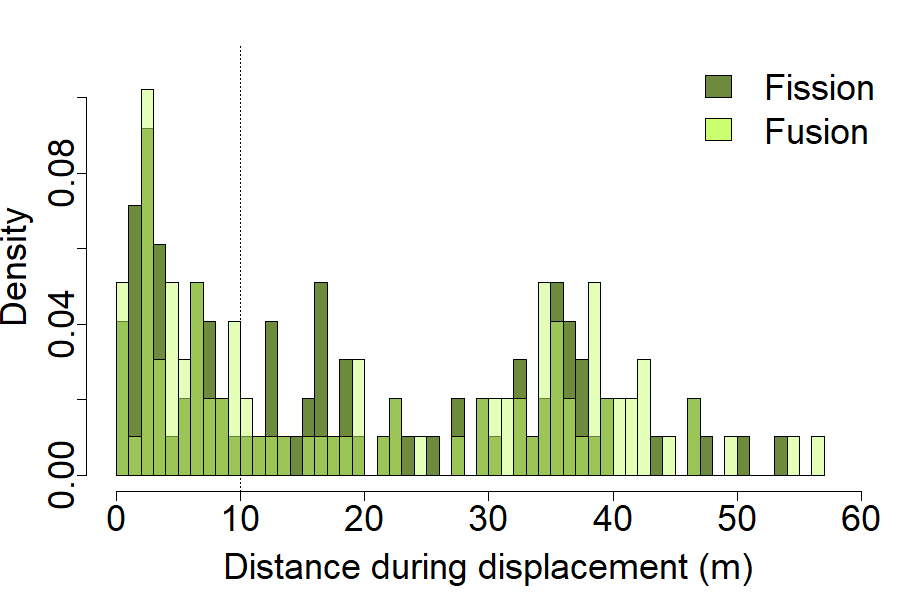

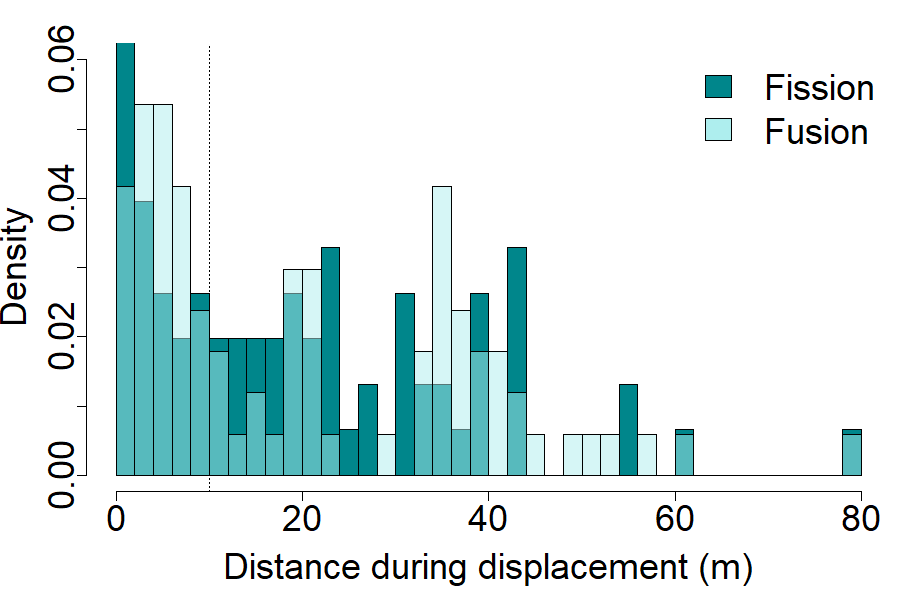

Figure S3. Distance distributions during fission and fusion events for Galaxy group (left), and Presidente group (right).

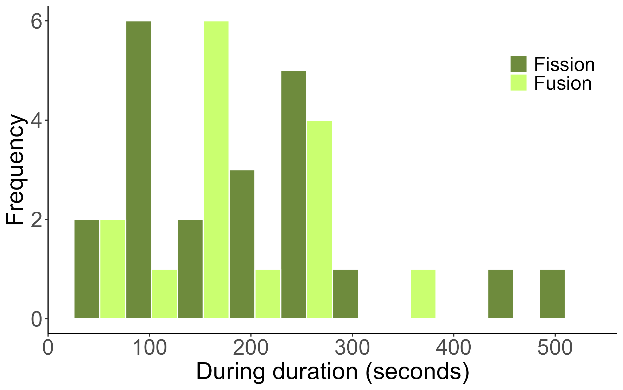

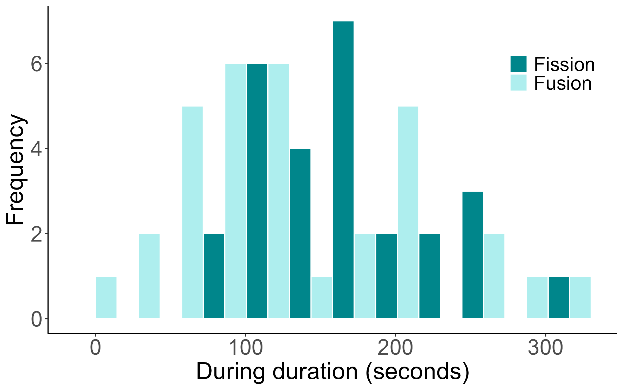

Figure S4. Distribution of the duration of the period during fission and fusion events for the Galaxy group (left) and the Presidente group (right).

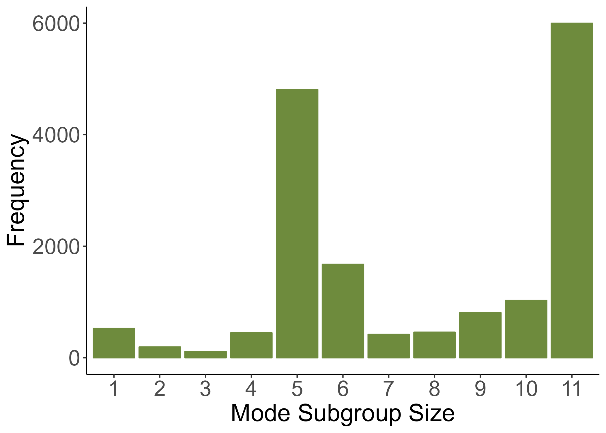

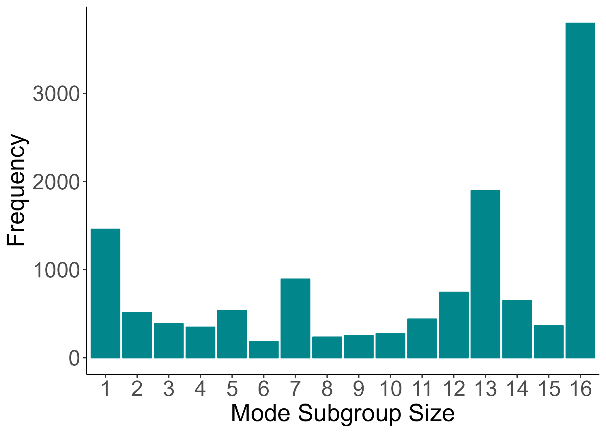

Figure S5. Distribution of subgroup sizes in the 1Hz period for the Galaxy group (left) and the Presidente group (right).

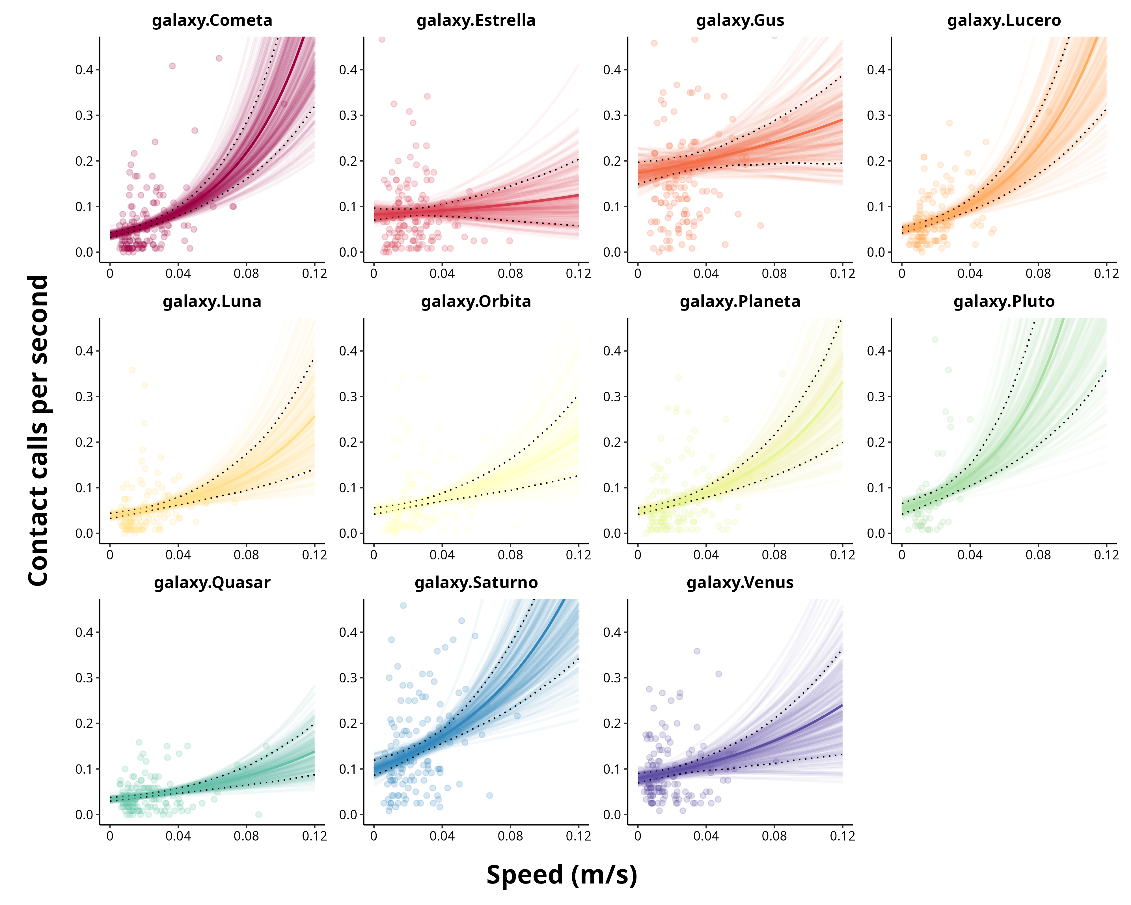

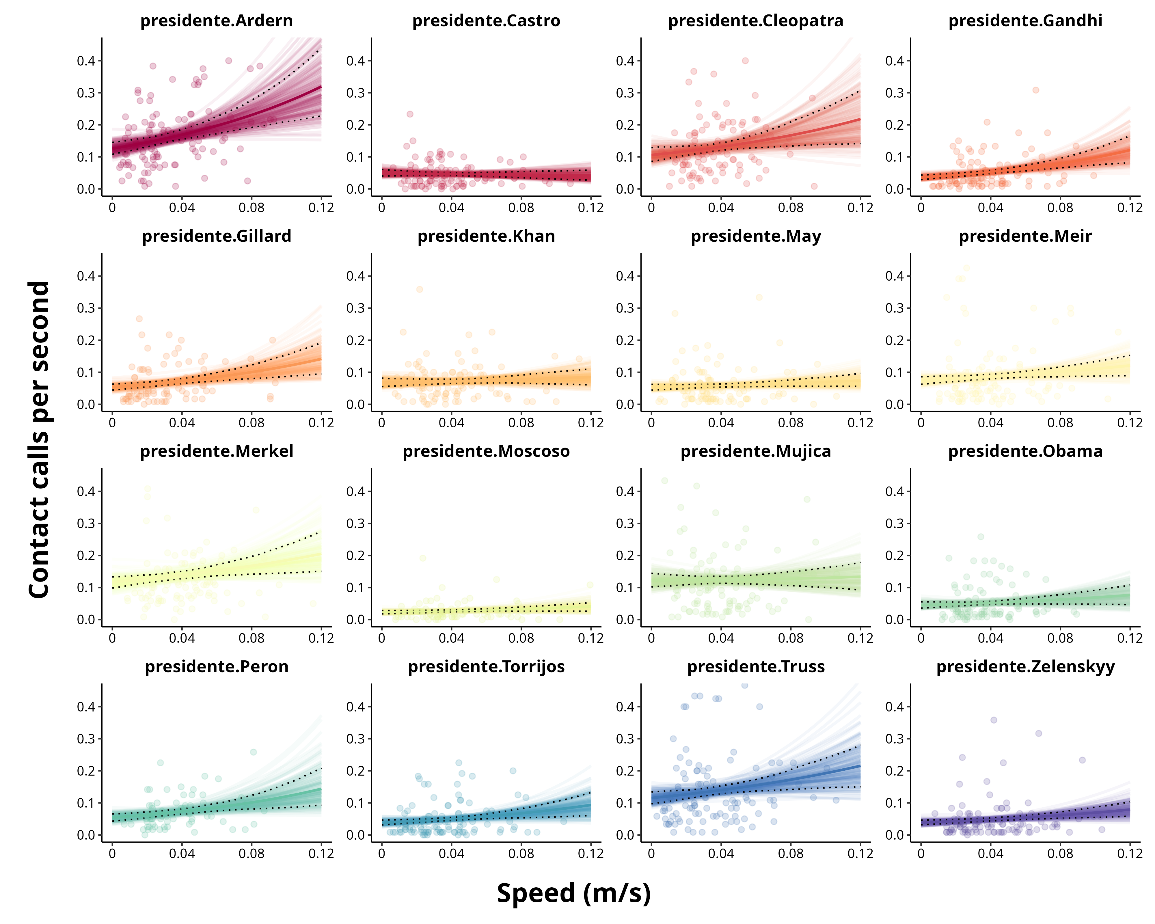

**B**

**A**

Figure S6. Model predictions for contact call rate as a function of travel speed for the Galaxy group (A) and the Presidente group (B). All panels show per-individual predictions, each represented by a different colour. Lighter lines are 100 randomly sampled posterior predictions, with the posterior median as a dark line. Black dotted lines show the 89% credible interval. Points represent raw data.

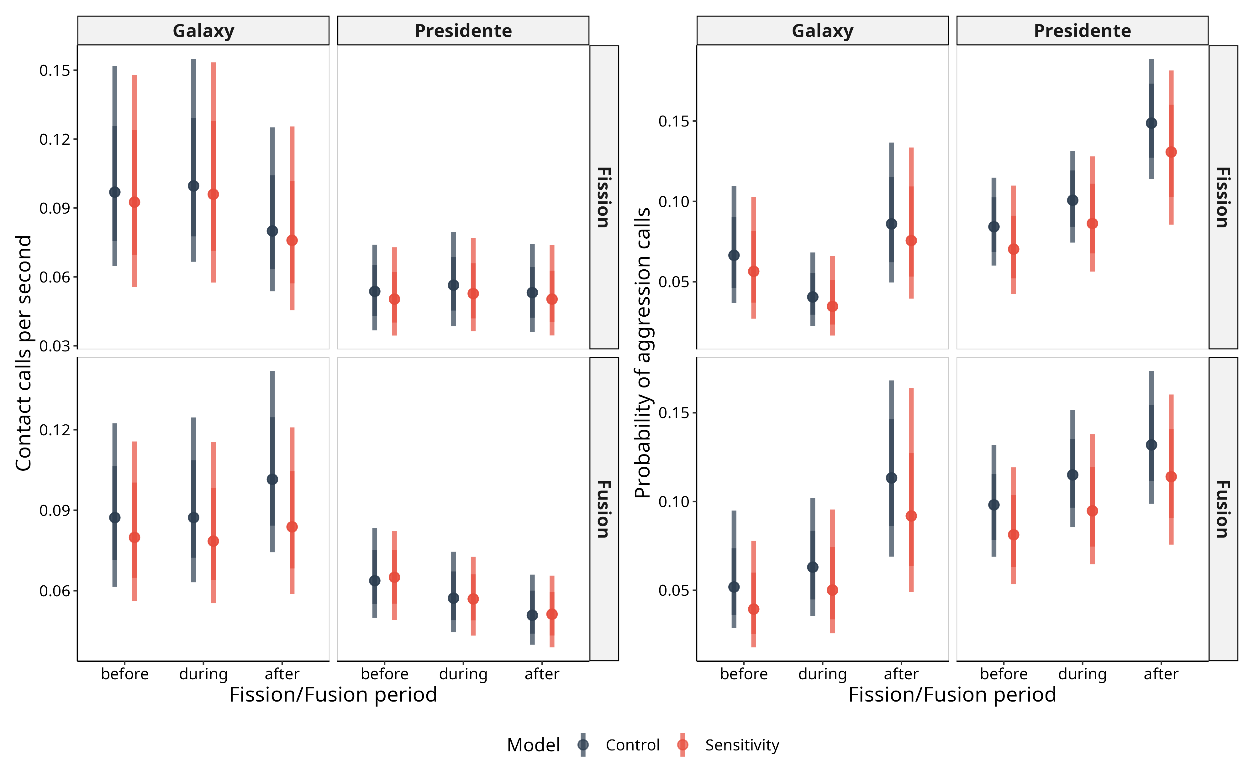

Figure S7. Sensitivity analysis assessing the effect of event-level non-independence on model estimates. Posterior predicted contact call rates (left) and probability of aggression calls (right) across fission and fusion event periods (before, during, after), shown separately for the Galaxy and Presidente groups. Control models include individual-level random intercepts only; sensitivity models include both individual-level and event-level random intercepts to account for potential non-independence between individuals within the same event. Both model types exclude baseline observations. Credible intervals represent the 66% (dark) and 89% (shaded) highest posterior density intervals.

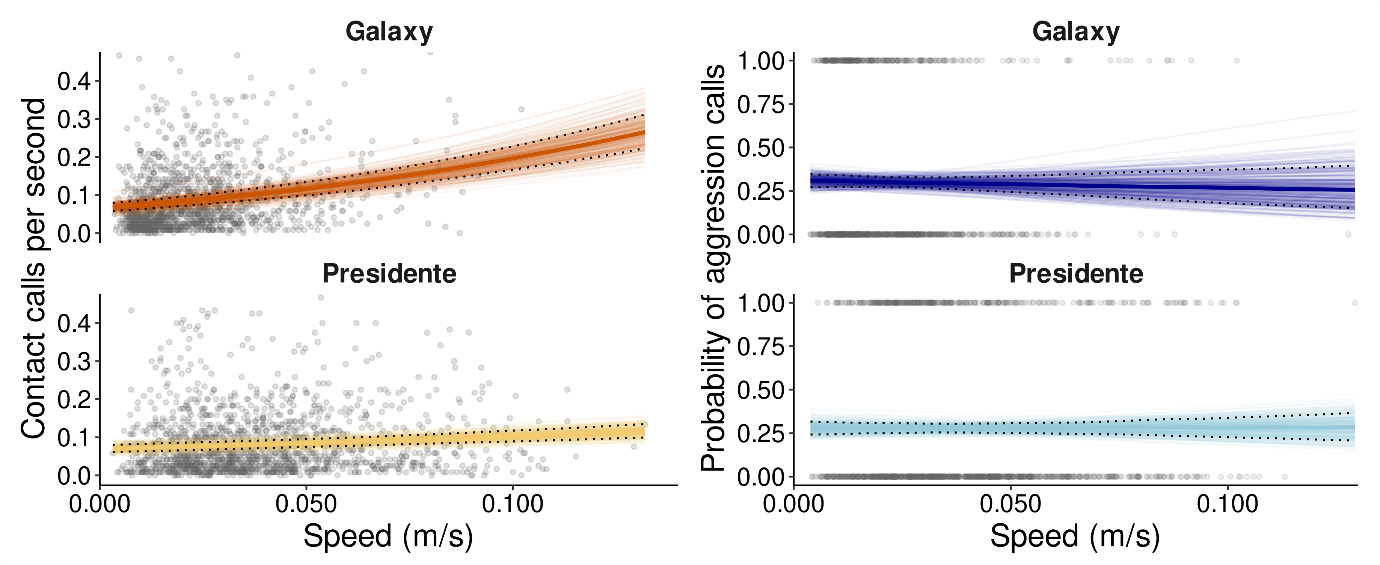
Figure S8. Model predictions for contact call rate (left) and probability of aggression calls (right) as a function of travel speed for the Galaxy group and the Presidente group. Lighter lines are 100 randomly sampled posterior predictions, with the posterior median as a dark line. Black dotted lines show the 89% credible interval. Grey points represent the raw data for each individual.
